# Saccharibacteria deploy two distinct Type IV pili, driving episymbiosis, host competition, and twitching motility

**DOI:** 10.1101/2024.11.25.624915

**Authors:** Alex S. Grossman, Lei Lei, Jack M. Botting, Jett Liu, Nusrat Nahar, João Gabriel S. Souza, Jun Liu, Jeffrey S. McLean, Xuesong He, Batbileg Bor

## Abstract

All cultivated Patescibacteria, or CPR, exist as obligate episymbionts on other microbes. Despite being ubiquitous in mammals and environmentally, molecular mechanisms of host identification and binding amongst ultrasmall bacterial episymbionts are largely unknown. Type 4 pili (T4P) are well conserved in this group and predicted to facilitate symbiotic interactions. To test this, we targeted T4P pilin genes in Saccharibacteria *Nanosynbacter lyticus* strain TM7x to assess their essentiality and roles in symbiosis. Our results revealed that *N. lyticus* assembles two distinct T4P, a non-essential thin pili that has the smallest diameter of any T4P and contributes to host-binding, episymbiont growth, and competitive fitness relative to other Saccharibacteria, and an essential thick pili whose functions include twitching motility. Identification of lectin-like minor pilins and modification of host cell walls suggest glycan binding mechanisms. Collectively our findings demonstrate that Saccharibacteria encode unique extracellular pili that are vital mediators of their underexplored episymbiotic lifestyle.

---

The bacteria of the superphylum Patescibacteria, historically known as the <u>c</u>andidate <u>p</u>hylum radiation or CPR, have long been considered part of the “microbial dark matter” due to their ubiquity in diverse metagenomic datasets, and the difficulty of stably culturing strains^1–3^. The ≈73 constituent phyla^4,5^ represent a monophyletic lineage hypothesized to contain 25%^6,7^ to 50%^8^ of total bacterial diversity based on phylogenetic analysis^9^. These bacteria feature ultrasmall sizes and highly reduced genomes with limited biosynthetic capacity typical of obligate symbionts^4,10^. Because most of these bacteria are predicted, or shown, to be dependent on other microorganisms, they cannot be isolated without a host organism. Experimental investigations into Patescibacteria have used co-isolation or enrichment methods to identify growth on the surfaces of bacteria^11^, surfaces of archaea^12^, or cytoplasmically within a paramecium^13^. All *in vitro* cultivation experiments have focused on Patescibacteria associated with bacterial host cells, specifically those within Saccharibacteria (historically known as TM7) that grow as epibionts on the surface of Actinomycetales hosts^14–16^.

Saccharibacteria occupy many ecosystems including groundwater samples^4^, soil samples^17^, invertebrate microbiomes^16,18^, vertebrate gut microbiomes^19^, and vertebrate oral microbiomes^20^. They are of particular interest within the human microbiome due to their prevalence in the oral cavity, gastrointestinal tract, respiratory tract, skin, urinary tract, and even breast milk^21^. Many isolated Saccharibacteria strains are derived from the oral microbiome where they are prevalent within tongue dorsum, subgingival, and periodontal biofilms^19,22,23^. Saccharibacteria have a long history of adaptation to oral ecosystems, evidenced by their DNA being detectable in mineralized dental calculus^24^ from medieval human remains^22,25^, neanderthal remains^22,26^, modern non-human primates ^27^, and even oral swabs from distantly related mammals, such as dolphins^19,28^. Investigation of microbiome transmission has demonstrated that Saccharibacteria are some of the most consistently transmitted oral microbes between mothers and infants^29^.

When growing episymbiotically on other bacteria, Saccharibacteria can profoundly alter the lifestyle and phenotype of their hosts. They can function as parasites causing a significant host death, they can induce changes in cell size and metabolic activity, and they can even provide advantages to a host population by protecting against bacteriophages^15,30–33^. When not attached to a host, Saccharibacteria are seemingly incapable of replication, however they are not metabolically inert. Studies have shown that catabolic pathways, like arginine deiminase systems, allow planktonic cells to maintain membrane integrity and infectivity during transitional life stages^34^. Despite the ubiquity of Saccharibacteria and their ability to alter host physiology, little is known about the molecular mechanisms underlying their metabolism and lifestyle. This dearth of information stems from their evolutionary divergence from non-Patescibacteria, difficulties associated with culturing, and, until recently, the lack of genetic tools^35^.

While Saccharibacteria encode multiple predicted surface proteins and structures capable of interacting with host surfaces, type <u>4 p</u>ili (henceforth T4P) are nearly universally encoded across the phylum^36^. T4P are complex polymeric filaments composed of pilin monomers. Structural pilins, called major pilins, make up the bulk of each filament, while less numerous minor pilins are integrated, typically at the tip, where they play diverse molecular roles^37^. Multiple classes of T4P have been identified that are typically distinguished by their signal peptides, the size of their pili, and the machinery proteins required^37^. Classes include type A pili (T4aP), present in many genera, type B pili (T4bP), most common in *Enterobacteriaceae*, and Tight adherence pili, also known as Fimbrial low-molecular weight protein pili^37,38^. In archetypical T4aP, utilization of T4P requires several machinery proteins, which have been characterized molecularly and structurally^39–42^. PilB and PilT are ATPases responsible for extending and retracting T4P respectively, while PilC, PilM, PilN, and PilO act as structural components anchoring the complex into the cell membrane and wall.

Previous investigation of Saccharibacteria strain TM7i using a non-specific chemical inhibitor of PilB revealed that T4P were required for twitching motility^16^ and Tn-seq analysis of S. epibionticum revealed that some T4P genes were essential for survival^35^. To establish a direct link between Saccharibacteria T4P and their essential functions, such as motility and host bacterial interaction, we used *Nanosynbacter lyticus* strain TM7x and its cognate host *Schaalia odontolytica* strain XH001 as a model system^11^. By adapting recently developed Saccharibacteria genetic manipulation protocols we systematically investigated each T4P pilin, their essentiality, their conservation, and their roles in bacterial host interaction, twitching motility, and competency.

## Results

### TM7x has multiple T4P loci, encoding two distinct systems

We bioinformatically identified all pili genes within *N. lyticus* TM7x and compared them to previously characterized T4P. A database of diverse T4P systems (T4aP, T4bP, Com, and Tad)^43^ was used to BLAST search the TM7x genome^44,45^. This provided IDs for 9 genes, 1 pilin and two loci encoding T4aP machinery genes. Complementary examination using TXSScan-1.1.0^46^ (E-value cutoff of 1*10^-5^) identified 8 pilins and 12 assembly genes spread across 4 genomic loci, also homologous to T4aP systems (Figure 1A and Table S1). Machinery genes were more similar to non-Patescibacteria representatives than pilin genes were to non-Patescibacteria pilins. All unannotated genes within the loci were examined in the AlphaFold database^47,48^ for N-terminal α-helices indicative of T4P pilins, revealing 4 additional pilins that were missed by sequence similarity searches. Whenever possible, genes were assigned the name of their closest T4P homolog, however due to the low homology of the pilins, many were assigned arbitrary letters (Figure 1B).

**Figure 1.**
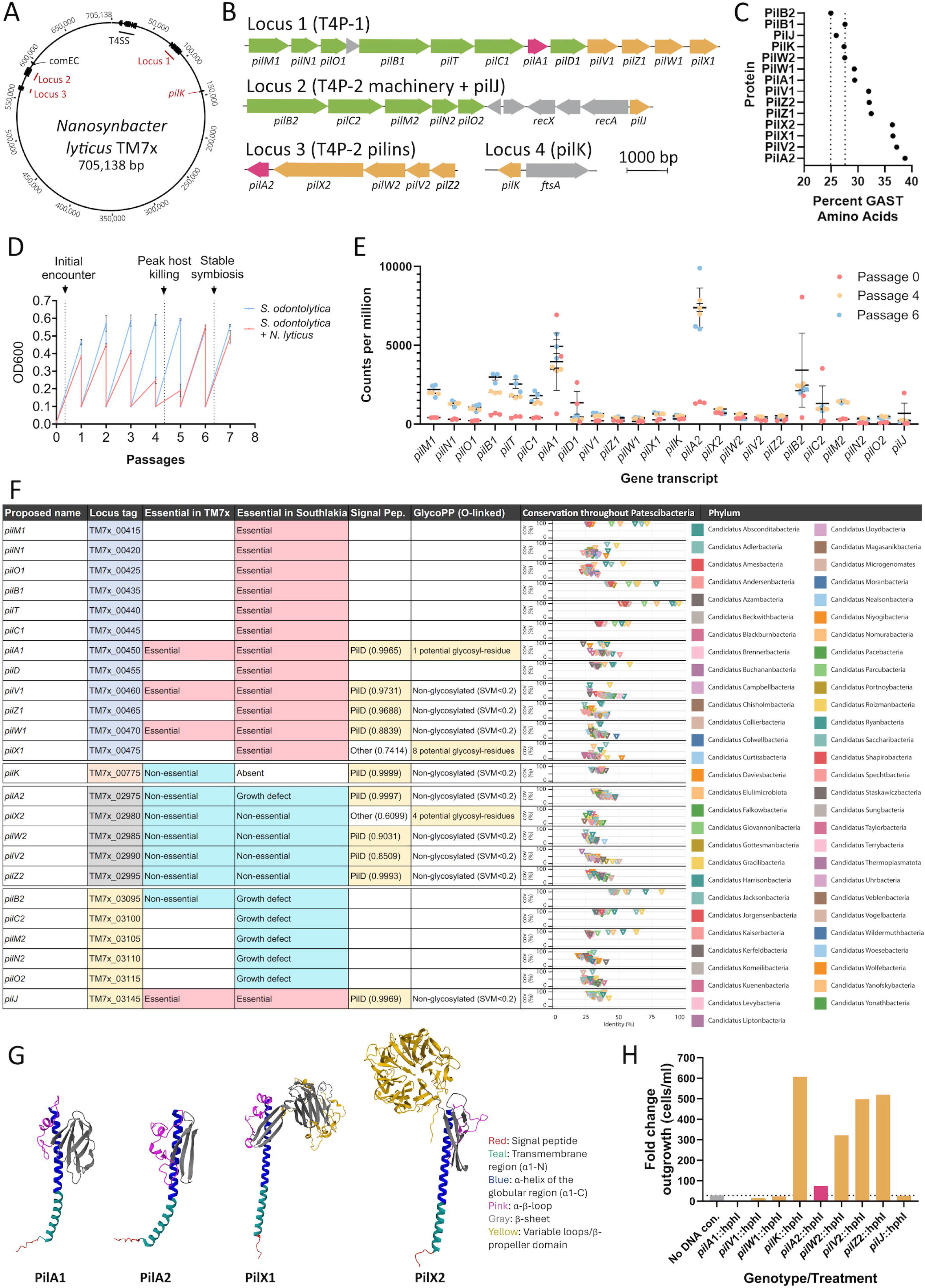
Identification and description of the T4P genes encoded by *N. lyticus*. Bioinformatic identification of T4P related genes within *N. lyticus* TM7x indicates that there are **(A)** four genomic loci containing T4P genes, which are not genomically associated with the T4SS. **(B)** These loci encode two distinct T4P extrusion systems, termed T4P-1 and T4P-2, as well as 12 putative pilins. Machinery genes are indicated in green, putative major pilin are in pink, and putative minor pilins are in yellow. **(C)** Examination of amino acid content indicated that pilin proteins, particularly those located in locus 3, contain a greater than average amount low-cost GAST residues. Transcriptomic data reported in Hendrickson *et al* 2022 examined expression of *N. lyticus* genes during initial host infection and stable association. **(D)** Passage growth dynamics are indicated for hosts in the presence (red) and absence (blue) of TM7x. **(E)** The majority of T4P genes were upregulated during symbiosis and exceptionally high expression of certain pilins (*pilA1* and *pilA2*) identifies major pilin subunits. **(F)** A table summarizes all *pil* genes named in this manuscript, compares their essentiality in *N. lyticus* to the related bacterium *S. epibioticum*, and indicates bioinformatically predicted modifications such as signal peptidase sites (with likelihood score) and potential for O-linked glycosylation (SVM threshold at 0.2). A database of genomes representing 56 Patescibacteria phyla was BLAST searched for homologs of each *pil* gene described. Graphs indicate the averaged percent coverage (y axes) vs percent identity (x axis). **(G)** AlphaFold2 structural predictions for the putative major pilins, as well as the most highly expressed minor pilins, reveal archetypical “lollipop” structures common to T4aP systems, with a partially hydrophobic N-terminal alpha helix followed by one or more hydrophilic globular domains. **(H)** Transformation with allelic exchange cassettes reveals which pilin genes are essential for *N. lyticus* survival and which are dispensable (*pilB2* and *pilX2* were mutated at a later time and not assessed) The outgrowth of mutant bacteria after transformation (cells per mL after final selection passage/cells per mL during final passage) also offers insight into potential growth defects introduced by loss of a gene.

T4P locus one (TM7x_00415-00475) encodes sufficient genes to produce a pilus, including the platform protein (PilC1), assembly ATPase (PilB1), retraction ATPase (PilT), alignment proteins (PilM1, PilN1, and PilO1), Signal Peptidase III (PilD), and 5 predicted pilins (Figures 1A and 1B). T4P locus two (TM7x_03095-03145) lacks a retraction ATPase but encodes a mostly complete set of assembly machinery (PilC2, PilB2, PilM2, PilN2, and PilO2), a recombinase operon (RecAX), and a single predicted pilin. T4P locus three (TM7x_02975-02995) encodes 5 predicted pilins. T4P locus four (TM7x_00775) contains an orphan pilin. Since T4P can be sheared from the cell surface mechanically, their constituent pilin proteins often have an increased abundance of low-cost amino acids like <u>G</u>ly, <u>A</u>la, <u>S</u>er, and <u>T</u>hr, collectively called GAST^49^. In *Pseudomonas aeruginosa*, major pilins have a GAST ≈38-47% while intracellular proteins have ≈24-32%^38^. Comparing GAST composition of predicted pilins to intracellular proteins (PilB1 and PilB2) revealed that this trend extends to TM7x pilins, particularly those encoded in loci one and three. Intracellular PilB sequences had 25.0-27.6% GAST. PilX1 (36.5%), PilX2 (36.4%), PilV2 (37.2%), and PilA2 (38.7%) had the highest GAST scores, indicating that these may be the most prone to loss via shearing (Figure 1C). PilW1, PilA1, PilV1, PilZ1, and PilZ2 had modestly elevated GAST scores (29.3%-32.5%). The orphan pilins PilJ and PilK, alongside PilW2 had GAST scores similar to intracellular proteins. Given the presence of redundant assembly machineries, we hypothesized that TM7x produces at least two distinct filaments: T4P-1 from loci one while T4P-2 from loci two.

To better understand *in vivo* function, we investigated existing transcriptomic datasets^32^ showing expression of TM7x genes during and after establishment of episymbiosis. When introduced into fresh host bacteria and subcultured/passaged to reach stability, TM7x undergo predictable growth dynamics (Figure 1D)^50^. Collectively, pilin expression accounted for ≈1.75% of TM7x transcripts, indicating that this is likely a metabolically expensive process. Expression of 21 out of 26 T4P genes was significantly lower during initial exposure to host bacteria (passage 0, 6 hours post-infection) than during peak host killing (passage 4), or during stable co-culture conditions (passage 6) (Figure 1E). The most highly expressed T4P genes during host association were *pilA2,* located in locus three, and *pilA1,* located in locus one, strongly suggesting that these are the major pilin subunits (Figure 1E). The most highly expressed putative minor pilin for T4P-1 was *pilX1*, which is also the largest pilin encoded in its locus. Similarly, the most highly expressed putative minor pilin within the same locus as *pilA2* was *pilX2*, which also carries a large accessory domain, demarcating these minor pilins as potential tip proteins.

Since T4P genes identified in *N. lyticus* TM7x had limited homology to T4P outside of Patescibacteria, we wanted to assess conservation of these genes within other Patescibacteria phyla. A database of 868 Patescibacteria genomes across 56 phyla^22^ (excluding Saccharibacteria) was assembled and BLAST analyzed for closest homologs to the identified T4P genes (Figure 1F and Table S2). Again, pilus assembly machinery genes were better conserved, with *pilB*, *pilT*, *pilM*, and *pilC* generally having homologs with 100% gene query coverage in most phyla at higher amino acid identities (25-75% identity). Conversely, pilin genes had very low levels of conservation, typically only sharing 10-25% coverage with query sequences. Since T4aP pilins share important N-terminal features (PilD signal peptides and transmembrane α-helices) the observed divergence suggests that their C-terminal, functional domains are likely diversifying.

To assess T4P conservation within Saccharibacteria, representative loci including close relatives of *N. lyticus* from *Nanosynbacteraceae*, *Saccharimonadaceae*, and GTDB f UBA1547 (clade G1) and distant relatives from *Nanosyncoccaceae* (G3), *Nanoperiomorbaceae* (G5), and *Nanogingivalaceae* (G6) were compared^22^ (Figure S1A). Amongst closely related strains, BB004, TM7-008, and *Southlakia epibionticum* ML1, all T4P associated genes were shared except *pilK* which only occurred in BB004 (Figures S1B-S1E).

More distant genomes within clade G1 (TM7i and *Nanosynbacter featherlites* BB001) were similar to TM7x, however each had lost one pilin encoded in locus three. Distant Saccharibacteria, *Nanoperiomorbus periodonticus* EAM2 (G5) and *Nanogingivalis gingiviticus* CMJM (G6), conserved T4P locus one, however had no homology to locus two, three, or four, suggesting that these organisms only produce a single T4P filament. Finally, the most distant Saccharibacteria *Nanosyncoccus nanoralicus* TM7-KMM (G3) encoded a T4P distinct in both synteny and sequence homology, suggesting an independent origin or divergent evolution.

Our results indicated that Saccharibacteria pilins were unique on a sequence level, however it was unclear how sequence dissimilarity translated into three-dimensional structures. AlphaFold2^51^ prediction of TM7x pilins demonstrated a range of diverse structures, particularly in the globular region (Figure S2A). The putative major pilins PilA1 and PilA2 have prepilin peptidase signal peptides (PilD) according to SignalP 6.0^52^ and are predicted to adopt classical “lollypop” structures associated with T4aP pilins^38,53,54^, wherein a globular domain is attached to an N-terminal hydrophobic α-helix (Figures 1G and S2). Putative minor pilins also have lollipop architecture, though most carry additional features on their globular domains (Figure S2A-B). Several are elongated by variable loops (PilV1, PilW1, PilV2, PilW2, and PilK) while others have additional C-terminal domains (PilX1 and PilX2). Specifically, PilX1 has a β-Jelly Roll-like domain absent from NCBI CD-Search^55^ and PilX2 has a heptabladed β-propeller domain with RCC1 repeats (Figure 1G). Interestingly, PilX1 and PilX2 were not predicted to have PilD signal peptides, potentially indicating that these pilins are post-translationally modified without PilD or prime pilus polymerization like PilY1 in *Myxococcus*^56^ (Figure 1F). Structural predictions for the extrusion/retraction machinery proteins PilB and PilT indicated that they were surprisingly similar to non-Patescibacteria T4aP, possessing well conserved Walker A boxes (ATP binding) and Walker B boxes (hydrolysis)^57^ (Figure S2C). Several previously discovered pilins, such as Neisserial PilE^58^, are post-translationally decorated with O-linked glycans that impact motility and virulence^59^. GlycoPP V2.0 predicted that three pilins are potentially O-glycosylated, PilA1, PilX1, and PilX2. PilA1 had a single site of potential glycosylation (Serine 88) while PilX1 and PilX2 had multiple sites of potential glycosylation^60,61^ (Figure 1F).

### Generation of T4P Pilin knockout mutants

We attempted to delete pilin genes from all four genomic loci, as well as extrusion ATPase *pilB2* via allelic exchange^35^. A neutral site mutant was generated (NS1::hphI) as a technical control. All targeted pilins in locus one (*pilA1*, *pilV1*, and *pilW1*) and two (*pilJ*) could not be deleted, suggesting that they are essential for TM7x survival and reproduction. All pilins in locus three (*pilA2*, *pilX2*, *pilW2*, *pilV2*, and *pilZ2*) and four (*pilK*) were non-essential (Figure 1H). The rate of TM7x population expansion post-selection was reduced in TM7xΔ*pilA2.* In Figure 1F we compare gene essentiality within TM7x to the related Saccharibacteria *S. epibionticum*^35^ and find that the results are in agreement, including the growth defect introduced by mutating *pilA2*. This suggests that transformation is consistently effective and that T4P may play similar roles within the families *Nanosynbacteraceae* and *Saccharimonadaceae*.

### Two distinct pili filaments are assembled on the surface of TM7x

To validate bioinformatic predictions, cryogenic-electron tomography (henceforth Cryo-ET) was used to observe TM7x surface filaments. Imaging predominantly focused on free floating TM7x^50^ cells. Tomograms of wildtype TM7x indicated ≈6.5 filaments/cells, with a bimodal distribution of diameters centered around 1.8 nm (2.5/cell) and 3.2 nm (4/cell), suggesting two distinct surface appendages (Figure 2A, 2B, 2K, 2L, and S3A-S3B), henceforth referred to as thin pili and thick pili. These widths are small for T4P, which typically have diameters between 5-8 nm^37^. Thinner than average T4P have been identified before, however TM7x’s thin pili appears to be the smallest yet described. Thin pili had an average length of 77.4 nm (range = 15-171 nm), and thick pili were significantly longer with an average length of 121.9 nm (range = 45-249 nm) (Figure 2M).

**Figure 2.**
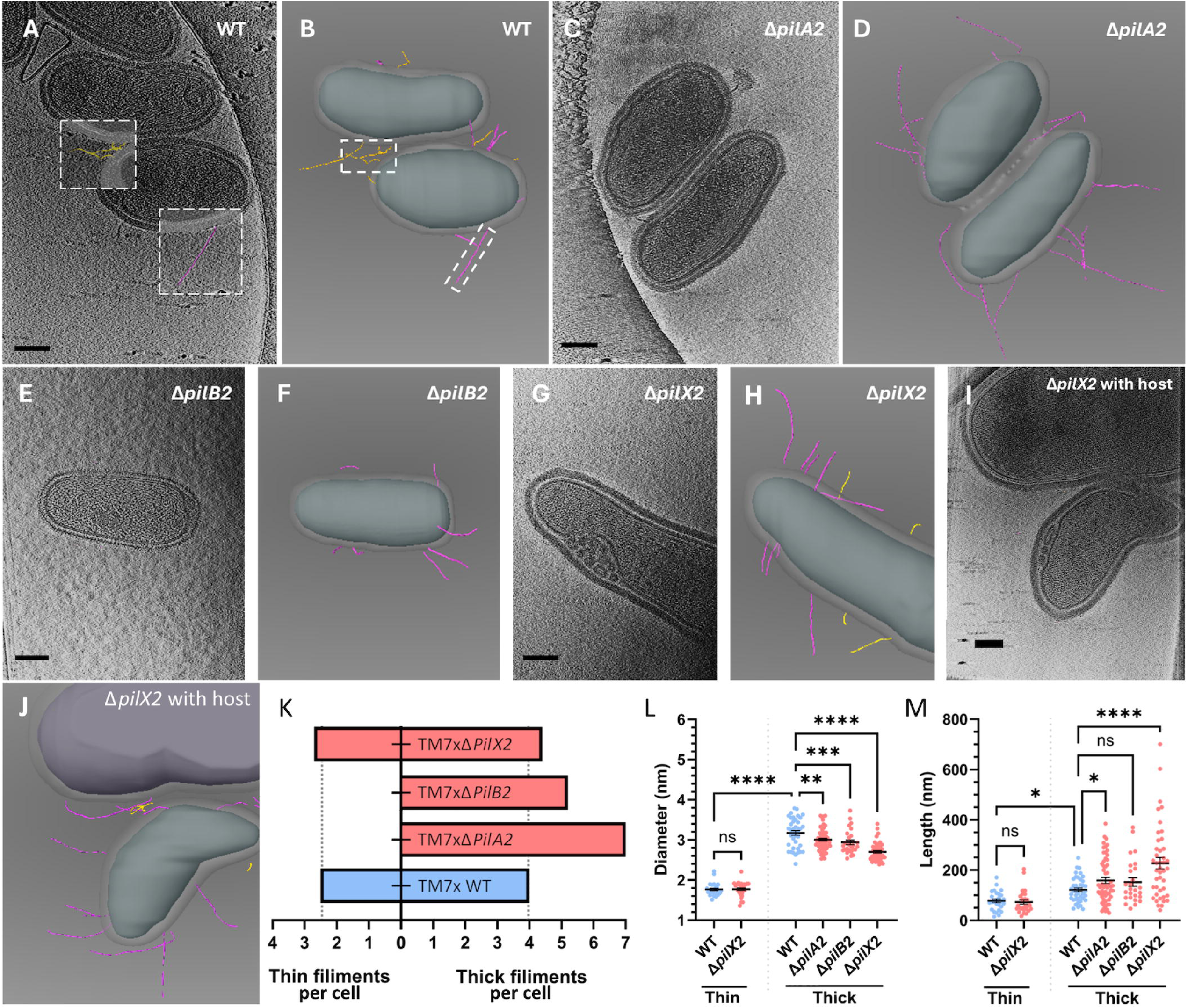
Cryo-ET reveals two distinct T4P filaments, distinguishable by their diameter. Electron tomography of planktonic wildtype *N. lyticus* TM7x **(A)** and subsequent 3D reconstruction of surface structures **(B)** reveals two distinct filament types (examples in white boxes) present on the surface, a thin pili with a diameter of ≈1.8 nm (highlighted in yellow) and a thick pili with a diameter of ≈3.2 nm (highlighted in purple). All scale bars are 100 nm long. Deletion of the putative major pilin PilA2 **(C-D)**, or the T4P-2 extrusion ATPase PilB2 **(E-F)** results in the loss of thin pili from cell surfaces, as well as modest changes in pili diameter and length **(K-M)**. Deletion of putative minor pilin PilX2 does not remove the thin filament from cells **(G-H)**, but it has a larger impact on the diameter and length of the thick filament. No mutant completely removed the ability for TM7x to associate with its host cells, as shown with a representative TM7xΔ*pilX2* **(IJ)**. * = p ≤ 0.05, ** = p ≤ 0.01, *** = p ≤ 0.001, **** = p ≤ 0.0001.

Next, mutants defective for the major pilin PilA2, the extrusion ATPase PilB2, and the minor pilin PilX2 were imaged. Tomograms of TM7xΔ*pilA2* (Figures 2C-2D and S3C-S3D) indicated ≈7 filaments/cell, composed entirely of thick pili (≈3.2 nm diameter) (Figures 2K and 2L). The thin filaments are completely removed by deleting PilA2, demonstrating that despite their narrow diameter the thin pili are T4P. Thick pili observed on TM7xΔ*pilA2* were more numerous (175% filaments/cell) and significantly longer (158.9 nm average) than wildtype thick pili (Figure 2M). Tomograms of TM7xΔ*pilB2* indicated ≈5.2 filaments/cell composed entirely of thick pili, much like TM7xΔ*pilA2* (Figures 2E-2F, 2K-2M, and S3E-S3F). Since loss of PilB2 removes thin pili like deletion of PilA2 does, we conclude that the thin pili are T4P-2, composed of assembly machinery encoded in locus two and pilins encoded in locus three. Increased thick filaments in both mutants suggests that loss of T4P-2 may result in a compensatory increase in T4P-1 expression. Finally, tomograms of TM7xΔ*pilX2* (Figures 2G-2H and S3G-S3H) indicate ≈4.4 thick filaments/cell and ≈2.7 thin filaments/cell (Figure 2K). While filament counts in TM7xΔ*pilX2* were similar to wildtype cells, identified thick filaments had significantly narrower diameters (2.7 nm average) and longer lengths (227.5 nm average) (Figures 2L and 2M). This shift suggests that either pilin subunit composition or conformational have been altered in these cells. Electron opacity differences between *S. odontolytica* and *N. lyticus* made it difficult to visualize host cells and TM7x simultaneously, however tomograms of TM7xΔ*pilX2* in contact with host cells show both filament types present but do not indicate concentration of either filament at their interface (Figures 2I-2J and S3G).

### T4P-2 contributes to TM7x host association, but not to motility or competency

T4P are important for many bacterial functions, including competency, twitching motility, adhesion, and assembly of bacterial communities^62^. First, we tested the contribution of T4P to episymbiotic host-binding, where axenic hosts were infected with each TM7x genotype at a <u>m</u>ultiplicity <u>o</u>f infection (MOI) of 0.1 (1:10 TM7x:host ratio), then percentage of hosts colonized by TM7x was quantified via phase-contrast microscopy. When exposed to wildtype TM7x for 24 hours, ≈10.6% of host cells bound TM7x (Figure 3A). Deletion of the minor pilins *pilK*, *pilW2*, *pilV2*, and *pilZ2* had no significant effect on host-binding. However, a significant defect in host-binding was induced by deletion of the major pilin *pilA2* (4.8% of host cells bound), the β-propeller bearing *pilX2* (4.6%), and the ATPase *pilB2* (3.9%) (Figure 3A). The similarity of the binding defects observed suggests that PilA2 and PilX2 are likely assembled into the same filaments and work in conjunction to interact with host bacteria. This is supported by the Cryo-ET results (Figure 2) where deletion of *pilA2* prevented assembly of T4P-2 (thin), and deletion of *pilX2* only impacted filament counts and length. Observed binding defects do not completely prevent host-binding, nor do they prevent the mutants from eventually reaching high densities (Figure 1H).

**Figure 3.**
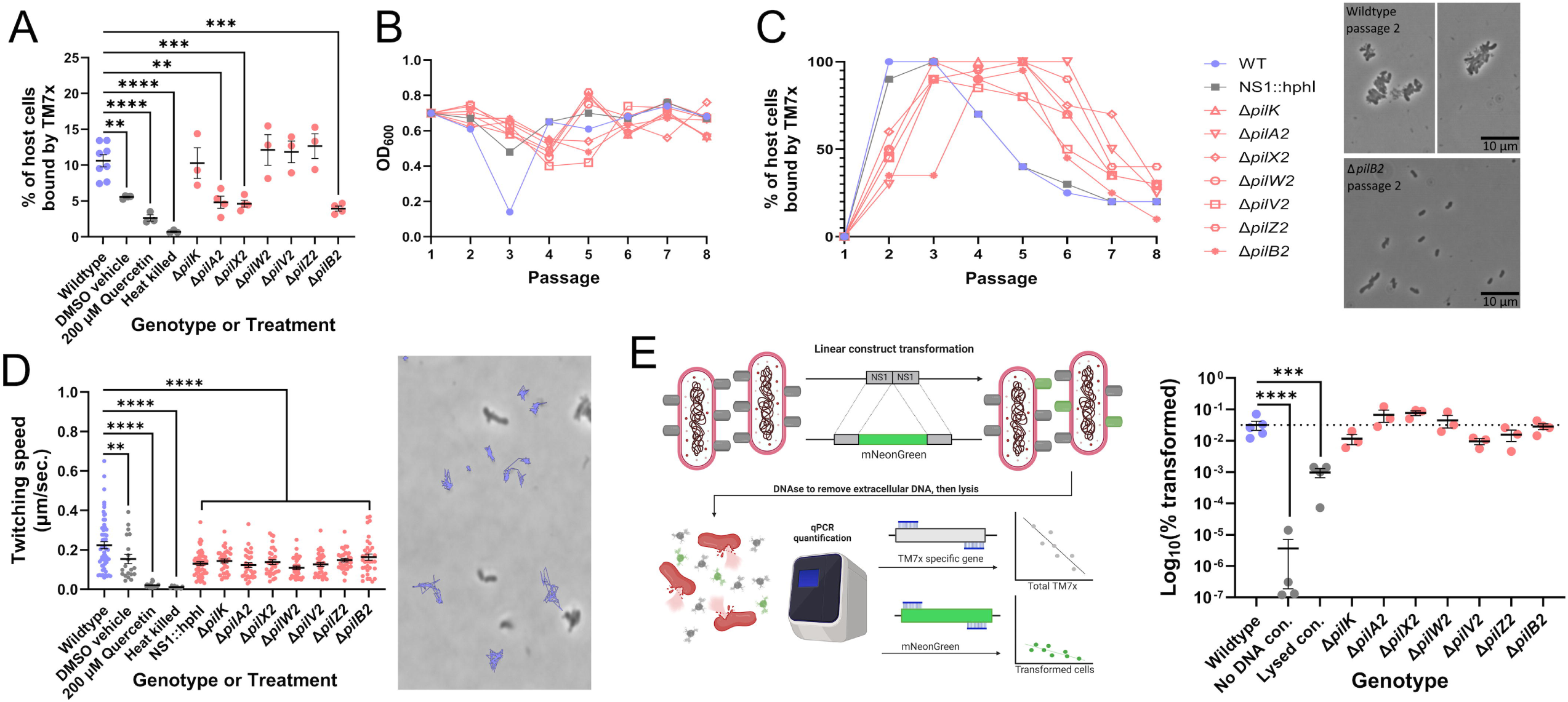
Characterization of *N. lyticus* TM7x pili gene mutants, examining common T4P roles such as host-binding, motility on surfaces, and genetic competence. Exposing *S. odontolytica* cultures to TM7x reveals ≈10.6% of hosts bound by wildtype episymbionts after 24 hours, which is reduced by approximately half in the PilA2, PilX2, and PilB2 mutants **(A)**. Application of non-specific PilB chemical inhibitor quercetin induces a similar reduction. Repeated passaging of episymbiont infected cultures reveals modestly reduced rates of host killing **(B)**, population growth, and host adaptation **(C)** in pilin mutants. Representative images of wildtype co-cultures and TM7xΔ*pilB2* co-cultures after 2 passages highlight the growth delay induced. Timelapse photography **(D)** of co-cultures allowed tracking of TM7x twitching motility and showed that all T4P mutants were motile, while the quercetin treated cells were not. A representative section of a timelapse image taken over 3 minutes demonstrates the stochastic twitching motions of wildtype TM7x. Competence assays **(E)** demonstrated that all pili gene mutants maintained wildtype levels of competence by transforming cells of each genotype with an mNeonGreen gene at a neutral site and using qPCR to quantify the percentage of cells successfully transformed. * = p ≤ 0.05, ** = p ≤ 0.01, *** = p ≤ 0.001, **** = p ≤ 0.0001.

To determine if pilin mutants have altered growth dynamics, or altered host growth dynamics, all mutants were isolated, re-introduced to axenic *S. odontolytica* cultures, and passaged daily until populations stabilized. At every passage OD_600_ was recorded, largely contributed by the host, and average percentage of host cells bound by TM7x was determined via microscopy. Wildtype TM7x induced host population crash at passage 3 as observed previously^50^, the neutral site mutant (NS1::hphI) had a smaller host crash at passage 3, and all pilin mutants induced a small host crash at passage 4 (Figure 3B). The reduced crash phenotype seen in all mutants suggests that the transformation process or cassette decreases the amount of stress TM7x applies to hosts. The delayed host die-off seen in T4P-2 mutants relative to the neutral mutant suggests that T4P-2 defects subtly impaired in host-binding or growth rate. Wildtype TM7x and the neutral mutant achieved high levels of host-binding by passage 2 (90-100%), most pilin mutants achieved high levels at passage 3, and TM7xΔ*pilB2* did not achieve similar levels until passage 4 (Figure 3C). The neutral mutant stabilized their population at passage 6 (20-30%) while T4P-2 mutants stabilized at passage 7. Collectively these data suggest that disruption of T4P-2 decreases host-binding rate, particularly during the first 24 hours.

T4P are known to power both gliding and twitching motility, thus all mutants were examined for motility defects^63^. When suspended on agar gel, Saccharibacteria can be seen to “twitch” rapidly across the surface. Wildtype TM7x had an average speed of ≈0.22 μm/second and a max speed of 0.65 μm/second (Figure 3D). To determine if TM7x twitching was T4P dependent, we treated cells with the ATPase chemical inhibitor quercetin, a vehicle control, and lethal temperatures. DMSO did lower twitching motility slightly (average 0.15 μm/second), however quercetin (0.02 μm/second) and heat treatment (average 0.01 μm/second) almost completely ablated motility (Figure 3D). Examining T4P mutants showed similar twitching defects to the neutral site mutant (average 0.13 μm/second). These results suggest that the Hygromycin B cassette or transformation process causes a subtle twitching motility defect, but loss of T4P-2 does not. Since the T4P inhibitor quercetin inhibited twitching, and loss of T4P-2 does not, our data indicate that T4P-1 likely drives twitching motility, which is supported by the presence of a retraction ATPase (PilT) homolog within the essential T4P locus one.

T4P contribute to natural competence in many bacteria^64,65^. Since Saccharibacteria do not encode a complete nucleic acid biosynthesis pathway^66,67^ and multiple Saccharibacters have proven to be naturally competent^35^, all T4P-2 mutants were tested for competence. A linear construct was assembled to insert a Green Fluorescent Protein (mNeonGreen) into neutral site 1 (Figure 3E). Co-cultures of *S. odontolytica* with each TM7x genotype were transformed, DNase treated to remove extracellular DNA, and quantified via qPCR (total cells vs transformed). Lysed cells were included to control for removal of extracellular DNA. ≈0.01%-0.07% of wildtype TM7x were transformed (≈1:3000 cells), lysed and untransformed controls were significantly lower than wildtype. Pilin mutants showed similar competency rates to wildtype (0.01%-0.08%). After normalization, no mutant was significantly different from wildtype, suggesting that T4P-2 does not contribute to competency (Figure 3E).

### Host cell wall polysaccharides are important for TM7x binding

To determine if minor pilins from TM7x may bind glycans to facilitate host interactions as seen in other bacteria^68–71^, we isolated C-terminal domain structures from PilX1 and PilX2 (predicted by AlphaFold3^47^) and submitted them to DALI^72,73^ for structural similarity searching. The best PilX1 structural model was similar to diverse carbohydrate hydrolases, several lectins, and a variety of capsid proteins (Table S4). The best PilX2 structural model was similar to diverse carbohydrate hydrolases, several fucose binding lectins, DNA binding proteins, and peptidases. Specifically, the predicted PilX1 model resembled L-type lectins^74^ and PilX2 resembled PLL-type lectins^75^ (Figure 4A-4B). Additionally, PilX2 had structural similarity to a single T4P related protein, PilY1 from *P. aeruginosa*, known to bind integrin glycoproteins^76^ and regulate twitching motility^77^.

**Figure 4.**
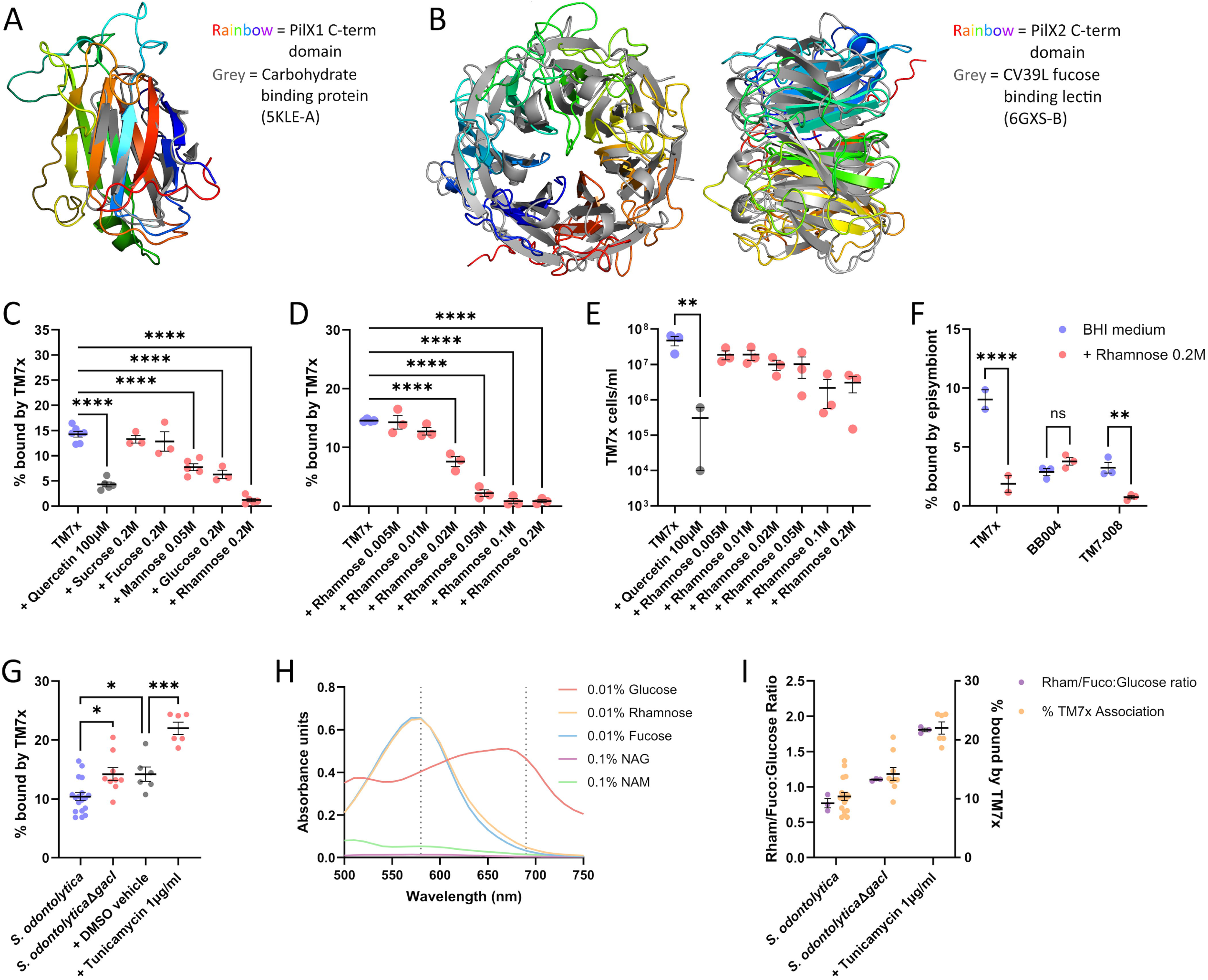
Exogenous sugars and cell wall glycans impact Saccharibacteria binding. DALI structural similarity searching revealed that both PilX1 and PilX2 have similarity to carbohydrate binding and cleaving proteins. Each pilin C-terminal domain (rainbow) was aligned to a high confidence (low RMSD, high z value) lectin-like structure (grey), **(A)** cellopentose binding protein CBM_E1 (5KLE-A) for PilX1 and **(B)** fucose binding lectin CV39L (6GXS-B) for PilX2. **(C)** Addition of exogenous sugars representing some of the most common cell wall glycosylation moieties indicates differential impact on TM7x bacterial host-binding. Soluble mannose and glucose approximately halved binding, and rhamnose almost completely inhibited binding. **(D)** A dosage-response curve of L-rhamnose reveals an IC50 value of 19.77 mM, despite total TM7x not being significantly decreased at any concentration **(E)**. Rhamnose-dependent binding inhibition is strain specific, appearing in co-cultures containing TM7x and TM7-008 but not in co-cultures containing BB004 **(F)**. Targeting genes associated with incorporation of Rha-CWPS into the cell wall shows that disruption of this pathway in *S. odontolytica* seemingly increases TM7x host-binding **(G)**. Specifically, loss of a putative N-acetylglucosyltransferase (GacI) and inhibition via tunicamycin increased binding. To investigate how these treatments impacted the concentrations of common cell wall glycans a modified anthrone assay was developed to quantify the ratio of rhamnose/fucose relative to glucose colorimetrically **(H)**. Combining these data **(I)** indicates a correlation between the rhamnose/fucose to glucose ratio and the exploitability of *S. odontolytica* as a host for TM7x. * = p ≤ 0.05, ** = p ≤ 0.01, *** = p ≤ 0.001, **** = p ≤ 0.0001.

To competitively inhibit potential lectin activity, TM7x was added to cultures of *S. odontolytica* supplemented with exogenous monosaccharides. Co-cultures were incubated with mannose, sucrose, fucose, glucose, or rhamnose for 24 hours. Sugar concentrations were optimized to avoid impacting host bacteria growth (Figure S4A). TM7x binding in the absence of sugars (≈14.3%) was not significantly different from sucrose (≈13.3%) or fucose (≈12.8%), however binding was significantly decreased with mannose (≈7.7%), glucose (≈6.3%), and, most drastically, rhamnose (≈1.2%) (Figure 4C). Quercetin significantly decreased binding (≈4.3%), but not as much as rhamnose. A rhamnose concentration response curve was generated between 200 mM and 5 mM, measuring frequency of host-binding (Figure 4D) and total TM7x (Figure 4E). Concentration dependent inhibition of host-binding was observed with an inhibition coefficient (IC50) of 19.77 mM (Figures S4B-S4C), despite there being no significant decrease in total TM7x. To determine if rhamnose-dependent binding inhibition was species specific or general to Saccharibacteria that encode T4P-2, three strains of *Nanosynbacteraceae* (TM7x (HMT-952), BB004 (HMT-488), and TM7-008 (HMT-352)) were grown in exogenous rhamnose. Interestingly, TM7-008 displayed decreased binding like TM7x, while there was no significant impact on BB004 binding (Figure 4F).

The native host organism of TM7x encodes homologs of rhamnan or rhamnose rich-cell wall polysaccharide (henceforth Rha-CWPS) biosynthesis proteins which may be contributing rhamnose to their cell walls. Within Streptococcal species, Rha-CWPS act similarly to wall teichoic acids^78^, contribute to phage and host interactions^79^, and can be used to identify immunological serotypes^80^. Within *S. odontolytica*, these genes are found in 4 genomic loci (Figure S4D) and include dDTP-rhamnose synthesis (*rml* genes), a rhamnose-fucose epimerase, Rha-CWPS extension and export (*rgp* genes), up to seven predicted glycosyltransferases used for addition of chain decorations, and a system for phosphocholine decoration of teichoic acids based on analogy to *S. pneumoniae*^81,82^. Rha-CWPS are not well characterized within Actinobacteria and have only been experimentally described within the family *Microbacteriaceae*^83^, however BLAST analysis using *rgpB* and *rgpF* from *S. odontolytica* reveals homologs throughout *Schaalia* and *Actinomyces* as well. Linear allelic exchange constructs were designed to delete *rmlC-D* (APY09_RS00100), *rgpB* (APY09_RS05560), *rgpF* (APY09_RS05510), *tagO* (APY09_RS06605), a putative galactosyltransferase (APY09_RS05455), a putative N-acetyl glucosyltransferase (APY09_RS00090), and a convergently encoded neutral site (between APY09_RS05760 and APY09_RS05765). After transformation, only the N-acetyl glucosyltransferase (henceforth *gacI*) and the neutral site could be successfully mutated, indicating that all other genes were essential for survival.

Fortunately, TagO has a known chemical inhibitor, tunicamycin, which we utilized to examine this essential gene^80,84^. Growth curves were performed to determine an optimal concentration between 0.1 μM and 11.8 μM. All tested concentrations reduced *S. odontolytica* growth rate with or without TM7x (Figures S4E-S4F). TM7x synergistically inhibited host growth, completely preventing growth at concentrations above 5.9 μM tunicamycin. Microscopic examination revealed abundant TM7x at all concentrations, suggesting that the chemical is less cytotoxic to the episymbiont than the host (Figure S4G). A concentration of 1.2 μM (1 μg/mL) was chosen for downstream experiments to allow *S. odontolytica* growth while maximizing cell wall impact.

*S. odontolytica*Δ*gacI* and tunicamycin treated cells were grown with TM7x for determination of binding potential and without TM7x for cell wall composition analysis. When assessing TM7x binding, we found that the *gacI* mutant (≈14.2%) had increased binding relative to WT (≈10.4%), and the tunicamycin treatment (≈22.0%) had increased binding relative to a DMSO vehicle control (≈14.2%) (Figure 4G). Purified cell walls from each treatment were assessed for relative sugar concentrations using a modified anthrone assay^85^. This assay converts sugars into chromophores with differentiable absorption spectra, while having no effect on NAG/NAM residues from peptidoglycan (Figure 4H). As fucose is a stereoisomer of rhamnose, they cannot be distinguished by this assay. Wildtype *S. odontolytica* had a rhamnose/fucose to glucose ratio of ≈0.77, while the *gacI* mutant (1.10) and the tunicamycin treated cells (1.81) contained significantly more rhamnose/fucose. Comparing TM7x binding of these hosts to their relative rhamnose/fucose:glucose ratios shows a strong correlation (Figure 4I). These results indicate that deletion of *gacI* may expose glycans for TM7x binding by preventing decoration of Rha-CWPS, tunicamycin treatment may alter incorporation of teichoic acids/Rha-CWPS, and relative ratios of cell wall sugars, such as rhamnose, may make host bacteria more susceptible to TM7x binding.

### Saccharibacteria species competition is modulated by T4P

Since TM7x pilin mutants showed defects in initial host interaction and growth dynamics, we wanted to know if loss of T4P impaired competitive fitness against other Saccharibacteria. Competition would be expected in the oral cavity, which contains hundreds of microbial species, composed of 1%-21% Saccharibacteria^14,86,87^. There are no prior studies on Saccharibacteria competition. To establish an assay, the three species of *Nanosynbacteraceae* used previously (TM7x, BB004, and TM7-008) were combined to create tripartite and pairwise competitions. In tripartite competition, all species were added to a host culture, passaged every 24 hours, and enumerated via qPCR using strain-specific primers (Table S5). TM7x consistently competed poorly during passage 1, had a growth spike at passage 2, and then persisted around 1*10^4^ cells/mL (Figure 5A, S5A, and S5B). BB004 steadily declined each passage until reaching near extinction (≈100-10 cells/mL). TM7-008 rapidly increased in abundance during the first few passages and remained the dominant episymbiont throughout.

**Figure 5.**
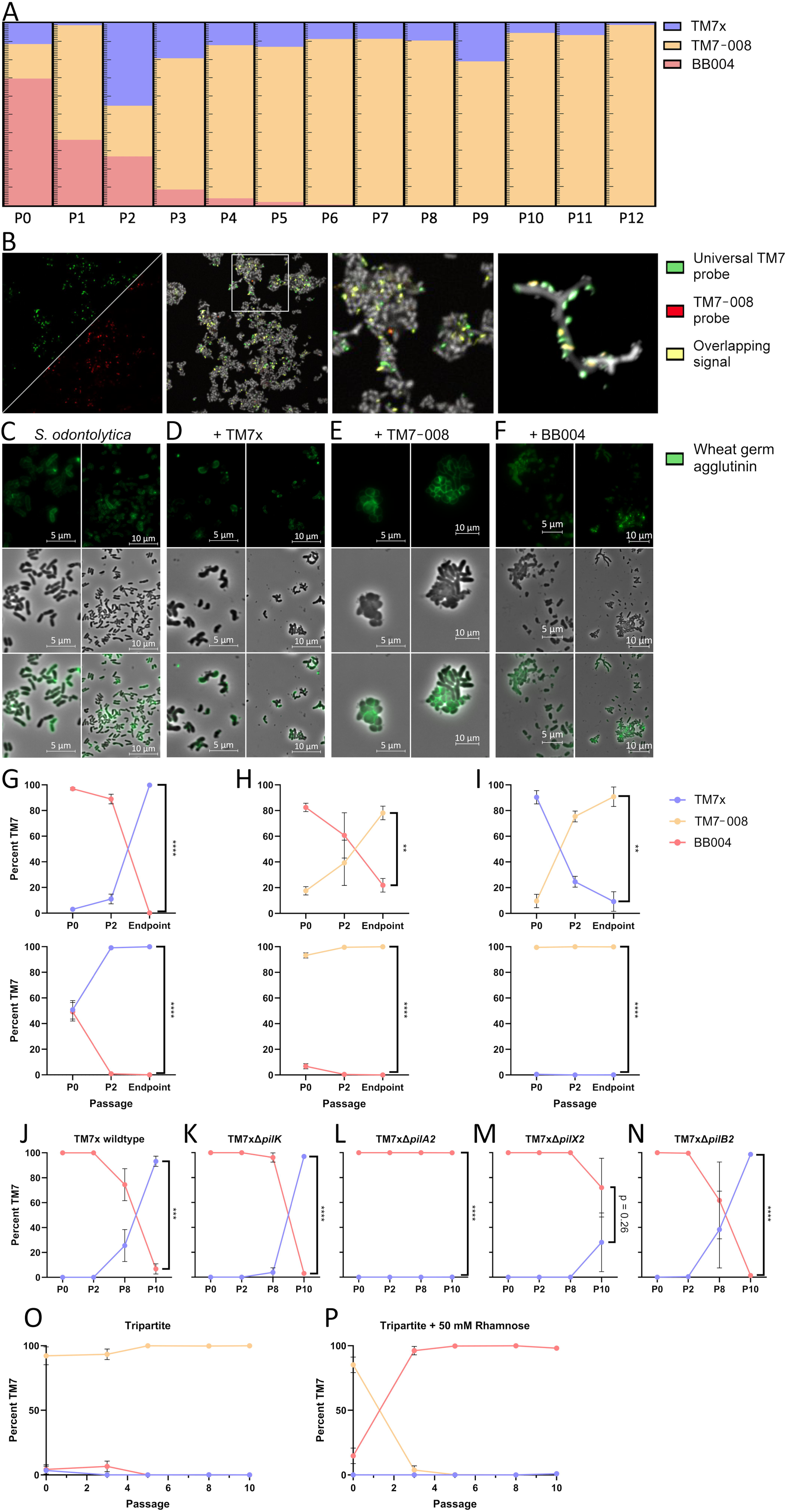
Saccharibacteria competition allows for host co-infection, results in consistent trophic successions, and is impacted by T4P-2. Co-infection of *S. odontolytica* cultures with three heterospecific Saccharibacteria strains (TM7x/blue, TM7-008/yellow, and BB004/red) followed by repeated passaging reveals consistent trophic successions, resulting in TM7-008 dominance **(A)**. Fluorescent in situ hybridization imaging using a general Saccharibacteria probe (green) and strain-specific probes (TM7-008 specific probe shown in red) highlights that competition strains can co-infect a single cell and do not show host or site exclusion **(B)**. Subsequent cell wall staining using conjugated WGA (green) reveals that Saccharibacteria strains have distinct effects on host morphology which could drive competitive outcomes, such as spheroid formation in TM7-008 co-cultures **(C-F)**. Pairwise combinations of TM7x/ BB004 **(G)**, TM7-008/ BB004 **(H)**, and TM7x/TM7-008 **(I)** reveal more consistent trophic successions, and demonstrate that while established host-episymbiont pairs have a temporal advantage in competition, priority effects are not enough to alter the outcome of competition/succession. By co-infecting bacterial host cells with TM7x T4P mutants and wildtype BB004 **(J-N)** we demonstrate that the host-association defects and/or growth defects induced in TM7xΔ*pilA2* and TM7xΔ*pilX2* significantly impair TM7x’s competitive fitness. Exogenous rhamnose increases competitive fitness of BB004 over strains with rhamnose-dependent binding inhibition **(O-P)**. TM7x and TM7-008. * = p ≤ 0.05, ** = p ≤ 0.01, *** = p ≤ 0.001, **** = p ≤ 0.0001. Absolute abundances and co-culture OD_600_ values in Figure S5.

To observe competition directly, we developed strain-specific Fluorescence *in situ* hybridization (FISH) probes. Three DNA probes were designed to target the 16S rRNA gene: Saccharibacteria universal probe (TM7-567)^88^, *N. lyticus* TM7x specific probe (TM7x-3), and TM7-008 specific probe (TM7-008-2) (Table S6). We designed multiple probes for BB004 but none of them were sufficiently specific due to shared sequence similarity with TM7x. To assess the specificity of TM7x-3 and TM7-008-2, cultures containing TM7x, BB004, or TM7-008 were multiplex stained with the universal Saccharibacteria probe and either TM7x-3 or TM7-008-2 (Figure S6A). Cultures of TM7x showed the greatest co-localization between TM7-567 and TM7x-3, indicating that TM7x-3 probe does not effectively hybridize to BB004 or TM7-008 (Figure S6B). Probe TM7-008-2 was similarly specific for TM7-008. TM7x vs TM7-008 competitions and tripartite competitions containing all three Saccharibacteria were multiplex stained at several time-points (Figure 5B and S7). At passage 2, Saccharibacteria existed diffusely throughout the population of host cells. When visualized again at passage 8, the density of Saccharibacteria had decreased, reflecting host-adaptation. When looking at single *S. odontolytica* cells, we can see TM7x and TM7-008 cells growing in close physical proximity on the same host (Figure 5B), showing no direct signs of spatial exclusion or neighbor killing.

Observation of *S. odontolytica* hosts within competition cultures indicated morphological shifts from pleomorphic bacilli to spheroids (Figure S7). To determine which microbe induced this shift, *S. odontolytica* cells were infected with each species individually, fixed, and stained with fluorophore conjugated <u>W</u>heat <u>G</u>erm <u>A</u>gglutinin lectin (WGA) to stain actively growing cell walls. Axenic *S. odontolytica* showed typical bacilli wherein cells had diffuse WGA binding with concentration at division septa. Cells infected with TM7x became irregular bacilli with reduced WGA incorporation (Figure 5C). WGA displayed patchy incorporation favoring polar regions, suggesting asymmetric cell wall remodeling (Figure 5D) consistent with previous studies that reported increased filamentation and branching in TM7x co-cultures^88^. Cells infected with TM7-008 adopted the spheroid morphology seen in tripartite competition cultures, suggesting that TM7-008’s ability to dominate competitions may result from induction of an altered host morphology or metabolic state (Figure 5E). Spheroid cells incorporated more WGA than bacilli in these cultures, indicating either inhibition of bacilli growth or increased WGA reactive epitope exposure. Infection by BB004 resulted in similar cell morphology and WGA incorporation to uninfected cells (Figure 5F), perhaps reflecting BB004’s poor competitive performance.

Next, we combined strains in a pairwise manner to determine their competitive hierarchy. To better represent an ecological succession of Saccharibacteria species, we introduced competitors into established episymbiotic co-cultures. Since *S. odontolytica* cells are known to adapt to TM7x association^89^, having an established co-culture could give a species a competitive advantage through priority effects^29,90^. When adding TM7x to established BB004, TM7x achieved dominance after passage 2, however when BB004 was added to established TM7x, TM7x became dominant before passage 2 (Figure 5G, S5C, and S5D). Having an established co-culture provided temporal advantage, but did not affect the outcome. When competing BB004 and TM7-008 a similar pattern emerged where TM7-008 always became dominant, but taking over an established co-culture required more than 2 passages (Figures 5H, S5E, and S5F). When competing TM7x and TM7-008, TM7-008 always became dominant and managed to do so by passage 2 (Figure 5I, S5G, and S5H). In summary, TM7-008 outcompeted both competitors, while TM7x only outcompeted BB004, illustrating that priority effects provided temporal advantage but did not impact ecological succession.

To provide an ecological context for TM7x T4P, TM7x’s ability to outcompete BB004 was examined using TM7xΔ*pilK*, TM7xΔ*pilA2*, TM7xΔ*pilX2,* and TM7xΔ*pilB2* mutants. When inoculated into these established BB004 co-cultures, TM7x became the predominant Saccharibacteria present between passage 8 and passage 10 (Figure 5J and S5I). TM7xΔ*pilK* and TM7xΔ*pilB2* behaved identically (Figures 5K, 5N, S5J, and S5L). Conversely, TM7xΔ*pilA2* and TM7xΔ*pilX2* grew more slowly and did not manage to outcompete BB004 by passage 10 (Figure 5L, 5M). Absolute abundances indicate that, while not becoming dominant, both mutants did start growing by passage 10 (Figure S5K and S5M). Like our binding and growth dynamic assays (Figures 3A-3B), this suggests that PilA2 and PilX2 mediate early bacterial host interactions important for Saccharibacteria competition, but are not essential for long term binding, which is likely mediated by more complex surface structures. In contrast, TM7xΔ*pilB2* disrupted binding and growth dynamics, but surprisingly did not have a significant competitive defect. Previous host-binding experiments showed that rhamnose-dependent binding inhibition was specific to TM7x and TM7-008, but not BB004 (Figure 3D). Consistent with this observation, tripartite competition in the presence of rhamnose reversed the competition hierarchy observed without sugar supplementation, with BB004 dominating co-cultures and TM7-008 nearly reaching extinction by passage 10 (Figures 5O-5P). Supplementation with rhamnose and loss of T4P-2 do not alter competitive outcomes identically, suggesting that rhamnose may confound multiple glycan binding interactions. Together these data suggest that initial interactions with a host are vital determinants of episymbiont success and may contribute to competition in the oral cavity.

## Discussion

Our investigation of T4P in Saccharibacteria reveals highly specialized and strongly conserved macromolecular filaments that contribute to cellular functions and host association. *N. lyticus* encodes two distinct T4P (T4P-1 and T4P-2). The identified pili share many structural features with canonical T4aP systems such as assembly mechanisms, GAST content, potential for glycosylation, and glycan binding. However, these pili are distinguished by their uniquely small diameters, their function in episymbiosis, and the presence of two evolutionarily distinct filaments. The presence of multiple filaments provides a potential division of labor for extracellular functions like twitching and host detection. The full extent of T4P-1’s functional roles cannot be determined at this time due to their essentiality for cell survival. This indicates that twitching motility is indispensable for TM7x or that T4P-1 is playing an as of yet undiscovered role. The presence of an essential pilin encoded adjacent to *recA* and the lack of competence defect in T4P mutants hints at DNA uptake. T4P-2 significantly increases the rate of host-binding and the competitive fitness of the epibiont. Based on this observation we propose that T4P-2 acts as an initial host recognition factor, wherein an adhesin molecule (potentially PilX2) binds a host receptor to anchor *N. lyticus*. Subsequent twitching motility through T4P-1 can bring anchored cells into physical contact so that additional close-proximity cell-surface adhesins can establish a direct bond, as previously proposed^35^. Multistep host recognition may be beneficial in a complex biofilm ecosystem such as those found within the human microbiome^91^.Binding experiments with exogenous sugars and *S. odontolytica* mutants revealed that cell wall glycans, particularly rhamnose, are crucial for TM7x binding and potentially promising receptor candidate. Finally, we demonstrate consistent competitive outcomes between Saccharibacteria growing episymbiotically on *S. odontolytica*, link those competitive outcomes to manipulation of host cell morphology, and demonstrate that competitive outcomes can be disrupted by mutation of the T4P-2 pilins PilA2 and PilX2, as well as by rhamnose.

The T4P produced by *N. lyticus* are distinct from canonical systems and reflect their unique episymbiotic lifestyle. While bacteria such as *T. thermophilus*^92^ and *V. parahaemolyticus*^93^ have been shown to produce two distinct T4P filaments, it is not a common arrangement. Expression of two metabolically expensive piliation systems can be difficult to support, especially for an organism with a 705 kb genome. Therefore, conservation of dual T4P throughout all clade G1 Saccharibacteria suggests that they provide significant fitness benefits to these ultrasmall episymbionts. The presence of two distinct systems indicates either a division of expression timing, which is not supported by our electron tomography or transcriptomics, or a division of otherwise incompatible functions. One division of labor suggested by our data could be retraction. T4P-1 encodes a retraction ATPase which could allow the pili to generate forces after binding a substrate, and in support of this we find that inhibition of T4P ATPases through quercetin stopped *N. lyticus* twitching motility. Conversely T4P-2 does not appear to encode a retraction ATPase and none of the mutants generated for this filament impacted twitching motility. Hypothetically, T4P-1 could be responsible for retraction dependent processes while T4P-2 acts as a more passive mediator of surface interactions within the oral environment. T4P utilization within the oral microbiome has not been extensively studied in the past, but many constituents encode T4P predicted to be involved in motility^94^ and the oral commensal and opportunistic pathogen *S. sanguinis* encodes a heteropolymeric T4P essential for both biofilm aggregation and internalization into endothelial cells^95^.

Another distinction between these T4P and canonical T4P is their size. With an average diameter of 1.8 nm, T4P-2 from TM7x appears to be the thinnest T4P yet described. T4P-1 (diameter of 3.2 nm) is also thinner than average, particularly for a retractile pilus implicated in twitching motility, which must apply forces after binding. For example, T4P from *M. xanthus*^96,97^ have been demonstrated to apply forces up to 149 pN using 6-7.5 nm filaments and T4P from *N. gonorrhoeae*^98,99^ can apply up to 110 pN with ≈6 nm filaments. Reduced pili widths seemingly reflect *N. lyticus*’ small cell size and may be a strategy for reducing the metabolic costs of piliation or may indicate that they do not need to routinely withstand large forces. The latter would support our hypothesized model where T4P-2 mediates initial host interactions. Notably, T4P-2 had a greater GAST composition than T4P-1, suggesting that it is somewhat expendable and more prone to loss from cells, either as a function of its narrow diameter or lack of a retraction ATPase. An interesting direction for future studies would be to measure the tensile strength of these filaments to determine if they have a similar width to strength ratio to other T4P or if they have evolved greater relative strength to compensate for reduced width. Another avenue would be determination of high-resolution structures of T4P-1 and T4P-2 via *in-situ* cryo-EM to examine the precise arrangement of pilins in these uniquely thin filaments.

*N. lyticus*, its close relatives, and their bacterial host *S. odontolytica* provide a model system for studying the previously unexplored competitive interactions between Patescibacteria. Low complexity episymbiont communities competing for bacterial hosts allow for fine-tuned systematic examination of specific strains and even proteins. Our results showed that competition between isolated Saccharibacteria resulted in consistent successions, wherein some strains were intrinsically stronger competitors regardless of priming effects, or differences in initial inoculum. Strains which competed better tended to be the ones that induced the largest phenotypic changes in their hosts (cell size, shape, branching, and WGA incorporation). These results suggest that competition is mediated predominantly by host manipulation strategies, and less by direct Saccharibacter-Saccharibacter interactions. This is supported by FISH results that show two different Saccharibacteria strains living proximally on a single host cell. However, additional studies are required to rule out the possibility of direct interactions and should include more diverse Saccharibacteria with more predatory/parasitic episymbiotic lifestyles, such as PM004 and AC001^67^. Interestingly, the major pilin PilA2 and the minor pilin PilX2 were essential for effective competition, however, no significant decrease was seen in the PilB2 mutant, despite having compromised host-binding and lacking visible thin filaments. These data indicate that T4P-2 improves competitive fitness, and that pili produced without PilX2 do not restore function. The lack of competitive defect for the PilB2 mutant might imply leaky extrusion of T4P-2 pilins through the T4P-1 machinery, however further research is needed to determine the fate of T4P-2 pilins in the absence of PilB2.

The T4P encoded by Saccharibacteria adapt previously identified T4P roles like twitching motility^92,100^, host binding^68,101,102^, multicellular community/biofilm formation^93^, and bacterial predation^71^ to life at a new, smaller scale. Our studies expand the range of T4P-dependent symbiosis to include episymbiotic bacteria-bacteria interactions. However, if T4P-2 is genuinely non-retractile as suggested by our data, its function may be more analogous to curli fibers than to other T4P, highlighting the versatility of these evolutionarily ancient structures. Within *E. coli*, curli fibers are thin amyloid surface filaments that non-retractably adhere surfaces to facilitate binding and biofilm formation^91^.

The dearth of previous studies examining T4P in Patescibacteria means that many open questions surround the assembly machinery superstructure, extrusion/retraction regulation, filament polymerization, and minor pilin integration. No single study could answer all these disparate questions, but our findings provide a foundation to elaborate upon. Both T4P systems described encoded a relatively small major pilin subunit (PilA1 and PilA2) which was highly expressed in transcriptomics, alongside a host of minor pilins with more elaborate architectures. Bioinformatic investigation of potential O-linked glycosylation revealed three pilins with one or more potential glycosylation sites, one major pilin, PilA1, as well as two minor pilins, PilX1 and PilX2. Glycosylation of pilins is not uncommon and can allow other bacteria to evade immune opsonization^103^ and phage recognition^104^. Future studies should attempt to confirm pilin glycosylation to learn more about the unexplored glycosylation processes in Saccharibacteria and their potential contribution to immune evasion within an epithelial environment.

For many of the T4P-2 minor pilin mutants (*pilW2*, *pilV2*, *pilZ2*, and maybe *PilK*), no detectable phenotype could be found. Several of the minor pilins identified had sequence level homology to T2SS pseudopilins (*pilV1*, *pilZ1*, *pilK*, *pilW2*, and *pilJ*), a feature previously linked to core minor pilins^105^ that function to prime the T4SS assembly machinery, precipitating polymerization of the major pilins and eventually becoming the tip complex after extrusion. Our results suggest that, except for PilX2, loss of any single minor pilin was not sufficient to impair T4P-2 function, either because they did not contribute to the functions assayed, the remaining minor pilins were sufficient, or the paralogous minor pilins from T4P-1 could perform the role instead. Furthermore, the presence of T4P-2 on the surface of TM7xΔ*pilX2* cells in electron tomography indicates that despite being required for efficient host-binding, PilX2 is not required for filament extrusion. These observations, alongside PilX2’s C-terminal β-propeller, non-canonical signal peptide, and potential for glycosylation make it an excellent candidate for the adhesin component of the thin T4P-2 filaments. Another T4P tip pilin possessing a C-terminal heptabladed β-propeller has been described previously in *P. aeruginosa*, called PilY1. PilY1 functions to bind eukaryotic integrin glycoproteins, as well as calcium, to facilitate twitching motility^76^. PilY1 did appear in our DALI similarity search, but with a relatively high root mean square distance (rmsd 5.0). This dissimilarity suggests that these proteins likely do not bind the same receptor but supports the idea that PilX2 is acting as an adhesin and may have specificity for a glycan or glycoprotein.

Our findings supported a distinct role for sugars, and specifically cell wall glycans, in *N. lyticus* host-binding. The addition of exogenous rhamnose strongly inhibited host association in a strain-specific manner, which is exciting since *S. odontolytica* encodes homologs of many genes utilized to synthesize rhamnan, also known as Rha-CWPS, in Streptococcal species^79,80,84^. Multiple genera within *Microbacteriaceae* have been demonstrated to synthesize species-specific Rha-CWPS (also called rhamnomannans), composed of rhamnose, mannose, glucose, galactose, and glucosamine^83,106,107^. Based on the glycopolymer synthesis loci encoded by *S. odontolytica* (Figure S4) and the nearly 1:1 rhamnose/fucose:glucose ratio observed, we propose that this organism is synthesizing Rha-CWPS, expanding the range known to produce these complex polysaccharides into the order *Actinomycetales*. We attempted to delete a range of genes to ablate incorporation of these large structural sugars, reduce their length, or even prevent them from being effectively decorated and found that most Rha-CWPS biosynthesis genes were essential, as they often are in other bacteria^79,80,84^. The essentiality of these genes suggests that, independent of their role in Saccharibacteria binding, Rha-CWPS are important cell wall components within *Actinomycetales*, an order renowned for specialized, pleomorphic, and hyphal cell morphologies.

One gene predicted to add glycosyl decoration of Rha-CWPS, *gacI*, was deleted and increased *N. lyticus* host-binding, suggesting that unmasked rhamnose may be a target. Unexpectedly, the introduction of tunicamycin increased the relative concentration of either rhamnose or fucose in cell walls. However, the increase in rhamnose/fucose again increased TM7x binding, highlighting their potential as a Saccharibacterial lectin target. Rha-CWPS are promising candidates for host identification because they can be quite structurally diverse due to complex branching glycan decorations^80,106,107^. Species or strain level diversity could allow specific binding of productive hosts, while avoiding non-productive binding to dead-end hosts.

TM7x host range is restricted to a single monophyletic clade of Schaalia strains, indicating phylogenetic preference^89^. Furthermore, observation that BB004 outcompetes other epibionts in the presence of excess rhamnose supports this hypothesis. The identification of specific lectins and specific glycan target moieties will require further investigation. One limitation of our study is that we were unable to directly demonstrate binding between pilin proteins and glycans or cell walls. While our studies highlight similar binding defects in T4P mutants and cultures supplemented with exogenous sugars, future studies will have to determine if sugar binding and T4P binding represent a single host-binding pathway, or two parallel means of host detection and binding.

Techniques and discoveries from our studies will facilitate further molecular level characterization of T4P in Saccharibacteria and throughout the many constituent phyla of Patescibacteria, while highlighting their importance to episymbiotic niches. T4P are versatile molecular machines that have evolved to fill many roles in distinct linages, and our findings expand on this evolutionary flexibility. By comparing T4P produced by Patescibacteria to those produced by non-Patescibacteria, we can identify features that have changed since the divergence of these ancient lineages and begin to characterize how episymbiotic bacteria have adapted their extracellular surfaces and appendages to live in intimate contact with other microbes. Furthermore, the lessons we learn about Patescibacteria host-binding factors, like T4P, may in turn help us predict things like episymbiont host range or optimal culture conditions for isolating organisms beyond Saccharibacteria.

## Materials and Methods

### Bioinformatics analysis

All CDS encoded by *N .lyticus* strain TM7x (GCF_000803625.1) were BLAST searched using the NCBI BLASTp webtool against well characterized T4aP, T4bP, Com, and Tad systems identified in Denise et al 2019^43^ to identify and isolate T4P loci. Subsequently, the genome was submitted to TXSScan-1.1.0^46^ using a lenient E-value cutoff of 1*10^-5^. Predicted structures for all genes contained within T4P loci were downloaded as PDB files from the AlphaFold2 database^47,48^ and examined for structural features typical for pilin genes (N-terminal hydrophobic α-helices and C-terminal β-sheets). Genes were assigned the name of their closest homolog identified in either dataset, many minor pilins were assigned arbitrary letter designations (Table S1). To perform GAST analysis, the percentage of Gly, Ala, Ser, and Thr encoded by each pilin was calculated and summed, then compared to the summed GAST value of known membrane anchored T4P proteins PilB1 and PilB2^49^. The transcriptomic data are derived from a dataset originally published in Hendrickson et al 2022 and were generated as previously described^32^. Reads per million values for all *pil* genes were extracted from the dataset and analyzed via one-way ANOVA with Dunnett’s multiple comparison.

A whole genome phylogeny of select Saccharibacteria species was performed as described previously^22^ with the following modifications. All genomes were analyzed using GTDB-Tk v2.4.0^108^ to identify homologous protein sets which were conserved across all assayed genomes, resulting in 104 homologous sets. Homolog sets were individually aligned using hmmalign 3.3.2^109^, and trimmed using BMGE (default settings with gap rate cut-off 0.1) to remove poorly aligned sections^110^. Maximum likelihood analysis was performed via IQtree (-st AA -m TEST -bb 1000 -alrt 1000)^111^ using 1,000 replicates and the results were visualized via Dendroscope v3.8.10^112^. IQTree’s ModelFinder Plus function^113^ was used to choose the General matrix substitution model^114^ with empirical frequencies and 4 gamma rates (LG+F+G4) by optimizing BIC. Nodes with less than 60% bootstrap support on the consensus tree were collapsed.

To examine potential syntenic orientation of T4P loci amongst Saccharibacteria, a subset of genomes from the phylogeny were BLASTed with the *pil* gene sequences identified in TM7x and their flanking regions. Where identified, syntenic loci were visualized in Geneious Prime 2024.0.7. If no homologs were found to either the *pil* genes or their flanking regions, the locus was assumed to be absent from that genome. To predict the signal peptides for T4P pilins, both PilFind v1.0^115^ and SignalP v6.0^52^ were consulted. SignalP predictions were chosen because they reported signal peptides for a larger number of pilins. Prediction of O-linked glycosylation was performed using the GlycoPP v2.0 webserver using a Composition profile of patterns (CPP) + Secondary Structure (SS) model^60,61^. The <u>S</u>upport <u>V</u>ector <u>M</u>achine (SVM) threshold of 0.2 was selected to increase the stringency of analysis and to exclude a low-confidence cluster of residues that had SVM values between 0.0 and 0.05. To examine conservation of identified T4P within the candidate phylum, genomes across Candidatus Patescibacteria were downloaded from NCBI (n=868) (as of Jan 2024). Protein sequences for the coding genes in these genomes were used to generate a custom BLAST database. Amino acid sequences from the 24 *pil* genes encoded by TM7x were then used as queries against this database. The percentage identity and query coverage were averaged at Phylum (NCBI derived sequences)/Class (GTDB derived sequences) level.

### Bacteriological culture conditions, quantification, and viability

Bacterial strain list describing sources and genotypes present in supplemental table 6. All *N. lyticus* and *S. odontolytica* broth cultures were grown in reduced <u>b</u>rain-<u>h</u>eart infusion medium (henceforth BHI) in a Whitley A35 workstation at 37°C, as were all other Saccharibacteria containing cultures. When grown on solid medium, cultures were plated on reduced BHI + 5% sheep’s blood. When not stated otherwise, cultures were grown in a microaerophilic gas mixture containing 2% O_2_, 5% CO_2_, and balanced with N_2_. During the initial passages of TM7x transformation cultures, cells were instead raised anaerobically in a gas mixture of 2.5% H_2_S, 10% CO_2_, and balanced with N_2_ in a Coy lab products Vinyl Anaerobic Chamber. Saccharibacteria were stored at −80°C in 15% Glycerol to maintain viability. To standardize the <u>m</u>ultiplicity <u>o</u>f infection (MOI) for competition experiments, strain-specific primers (Table S4) were designed to quantify TM7x (Cells/mL = 10^0.9767*log_10_*Ct*+10.85^), BB004 (Cells/mL = 10^-0.2951 * *CT* + 13.611^), and TM7-008 (Cells/mL = 10^-0.2893 * *CT* + 13.362^). For host-binding experiments, competence, and twitching motility Saccharibacteria stocks were directly enumerated using a Nanosight pro (Malvern Panalytical) allowing generation of a standard curve correlating OD_600_ to cells/mL (Cells/mL = 10^0.9767*log_10_(*CT*) + 10.85^). *S. odontolytica* growth curves treated with a range of tunicamycin concentrations (0.1 μM – 11.8 μM) were performed in biological triplicate under the above microaerophilic conditions using a Cerillo Stratus microplate reader.

### *Nanosynbacter lyticus* genetics

All primers used to generate transformation constructs, screen for successful transformants, sequence insertion sites, and quantify TM7x via qPCR are described in supplemental table 4. Linear constructs for insertion of a hygromycin B resistance cassette were graciously provided by the Mougous lab. To construct a hygromycin B cassette that is expressed in TM7x, the promoter region and terminator region from elongation factor T (pTuf) in *S. epibionticum* were PCR amplified alongside the *hph* gene from *S. hygroscopicus* and assembled into the HphI cassette via NEB HiFi Assembly. To develop our transformation protocol and generate a neutral mutant control, the TM7x genome was searched to identify convergently encoded intergenic regions which would be unlikely to cause deleterious effects if modified, leading to the utilization of neutral site 1, henceforth NS1 (C-terminal UTR of TM7x_01290, position 244,491). For all gene knockout constructs, as well as the neutral site insertion, PCR amplification was used to amplify 300 bp homology arms upstream and downstream of each insertion site. NEB HiFi assembly (E2621) followed by PCR amplification was then used to generate linear constructs for transformation into each site. For each transformation, a co-culture of TM7x and host cells were grown overnight anaerobically, then 500 μL of overnight culture were mixed with 1 μg of each linear DNA construct and incubated for 6 hours. Additionally, a no hygromycin B control and a no DNA control were included to monitor selection of TM7x during transformation. After incubation, 2.5 mL of dilute *S. odontolytica* (OD_600_ = 0.05 in BHI) was added to each culture to provide new hosts for outgrowth of TM7x and hygromycin B was added to a concentration of 150 μg/mL. All cultures were incubated anaerobically for 24 hours, subsampled to assess TM7x concentration via qPCR, and then passaged 1:10 into dilute *S. odontolytica* to allow outgrowth of transformed cells. Cultures were passaged in this manner 4 times. Passage 5 was a “recovery passage”, grown without hygromycin B and under microaerophilic conditions to enhance outgrowth. Then transformation cultures were passage 1-3 additional times microaerophilically to allow complete outgrowth of the transformed cells. Whenever the NS1 transformation culture reached 70-100% *N. lyticus* saturation, the cultures were serially diluted and plated on BHI + 5% sheep’s blood to grow isolated colonies. After incubation for 48-72 hours, plates were observed under a stereomicroscope to identify “irregular” colonies with non-circular shapes because this colony morphology results from TM7x association^50^. PCR amplification was used to screen transformants for the presence of the HphI cassette, (≈6.3% of colonies transformed when using NS1::HphI). All mutants were confirmed via whole genome sequencing performed by Plasmidsaurus using Oxford Nanopore Technology.

### Cryogenic-electron tomography

Cryo-ET samples were prepared using copper grids with holey carbon support film (300 mesh, R2/2, Quantifoil) as previously described^116^. The grids were glow-discharged for ≈30 seconds before depositing 5 μL of TM7x/host cell solution in PBS (OD_600_ ≈1.25) with added 10nm gold fiducial beads onto them. Then, the grids were blotted with Whatman filter paper from the back side for about 8 s and rapidly frozen in a cryogenic mixture of 37% liquid ethane and 63% liquid propane cooled with liquid nitrogen using a homemade gravity-driven plunger apparatus or a Leica GP2 plunger set to 25° and 95% relative humidity (Leica Microsystems, Wetzlar, Germany).

The plunge-frozen grids were clipped into AutoGrids and mounted into the autoloader under liquid nitrogen and docked onto a 300 kV Titan Krios electron microscope (Thermo Fisher Scientific) that was equipped with a K3 BioQuantum direct electron detector (Gatan), a Volta Phase Plate (VPP), and BioQuantum energy filter (Gatan). SerialEM^117^ was used to collect single-axis tilt series around −5 μm defocus with VPP, with a cumulative dose of ≈60 e^−^/Å covering tilt angles from −48° to 48° (3° increment). Images were acquired in a dose-symmetric scheme using a FastTomo script with an effective pixel size of 2.148 Å at the specimen level. All recorded images were first drift corrected by the software MotionCor2^118^ and then stacked by the software package IMOD^117,119^. The tilt series were then aligned by IMOD with the patching tracking method. Tomograms were then reconstructed in IMOD using the SIRT algorithm. Segmentations and 3D renderings were generated in Dragonfly, version 2022.2 (Comet Technologies Canada Inc., Montreal, Canada). Values for pili diameter and length were analyzed via one-way ANOVA with Dunnett’s multiple comparison.

### *Schaalia odontolytica* binding assay

Prior to re-association experiments, all *TM7x* strains had to be isolated as previously described^50^. Briefly, 100 - 300 mL co-cultures containing each genotype were grown overnight, spun gently to remove host cells, filtered via 0.45 μm PVDF filters (Millipore, S2HVU02RE), spun at 30,000xG for 30 minutes to pellet Saccharibacteria, and normalized to an OD_600_ of 0.1. Three to four isolations were performed for each genotype, except wildtype which was replicated with every experiment to account for variation (N=8). Isolated Saccharibacteria cells were stored at −80°C in 15% glycerol in PBS (w/v). Isolates were normalized by OD600 and re-introduced to *S. odontolytica* hosts at an MOI of 0.1. Introduced co-cultures were incubated for 24 hours, then percentage of host cells bound by episymbiont’s was determined via phase contrast microscopy using a Nikon Eclipse E400 using the SPOT Advanced v4.6 software. For every biological replicate, 8-10 microscope fields were counted manually, containing between 218 and 622 individual cells (389 average). The average percentage bound was compared via one-way ANOVA with Dunnett’s multiple comparison test. For binding experiments with exogenous sugars, custom batches of BHI were prepared with sucrose, fucose, mannose, glucose, or rhamnose at 0.2M, lower concentrations were achieved by diluting with standard BHI. For testing mannose, a maximum concentration of 0.05M was used because greater concentrations inhibited effective growth of host cells. qPCR using primers #145-146 was used to assess total TM7x presence in rhamnose treated binding assays. For quercetin treatments (100 μM) and tunicamycin (1.2 μM) treatments, 1000x stocks were prepared fresh in DMSO and added to cultures after inoculation.

### *N. lyticus* pilin mutant growth dynamics

Prior to re-association experiments, all *TM7x* strains had to be isolated as previously described^50^. Each TM7x genotype was added to *S. odontolytica* at an MOI of 0.1, and then incubated according to above method. Every 24 hours, OD_600_ (ThermoScientific Genesys30) and phase contrast images (Nikon Eclipse E400) would be acquired prior to cultures being passaged via 1:10 dilution into fresh BHI media.

### Saccharibacteria twitching motility

Overnight co-cultures of *S. odontolytica* and TM7x, as well as adapted co-cultures containing all TM7x pilin mutants, were placed onto ≈350 μm thick 0.8% agar pads (w/v) to limit mobility and allowed to dry for 5 minutes. Agar pads were inverted and placed onto MatTek glass-bottom petri dishes. Samples were imaged under anaerobic conditions using phase contrast microscopy on a Nikon ECLIPSE Ti using the NIS Elements 5.30.02 software. Images were taken every 30 seconds for 3 minutes and all genotypes were imaged at least twice. Paths of movement were traced and measured in Fiji^120^. Between 31 and 34 cells were traced for every tested genotype except WT (N=58) and Neutral site 1 (N=48). Wildtype *N. lyticus* were also treated with 200 μM quercetin and DMSO vehicle to test T4P (N=20). Finally, cells were heated for 10 minutes at 97°C as a heat-killed control (N=10). Average twitching speeds were compared using a one-way ANOVA with Dunnett’s multiple comparison.

### *N. lyticus* competence assay

All primers used to generate transformation constructs and quantify transformants via qPCR are described in supplemental table 4. Linear constructs were designed to insert a codon optimized pTuf mNeonGreen expression cassette into the neutral site previously used for hygromycin B resistance transformation (NS1). NEB HiFi assembly master mix (E2621) was utilized to incorporate 300 bp homology arms to the cassette and PCR was used to amplify the linear construct. Co-cultures containing each *N. lyticus* strain were grown overnight in biological triplicate, then 500 μL of each overnight co-culture was mixed with 1 μg of NS1::mNeonGreen construct and incubated for 6 hours. Additionally, no DNA controls and lysed cell controls were performed using wildtype *N. lyticus*. After incubation with the construct, 2.5 mL of dilute *S. odontolytica* (OD_600_ = 0.05 in BHI) was added to each culture to provide new hosts for outgrowth. After 24 hours of anerobic growth, cells were rinsed 2x in PBS. The lysed cells control was heated to 97°C for 10 min, frozen, and heated again for 5 minutes. All treatments were then treated with NEB DNase I (2U/mL, M0303) for 30 minutes to remove extracellular DNA. gDNA was extracted from each co-culture using MasterPure Gram-Positive DNA Purification kits (MGP04100). gDNA was diluted 1:10 and then qPCR using TM7x specific primers and mNeonGreen specific primers was used to assess the percentage of cells now carrying the transformation cassette. Data was log_10_ transformed prior to analysis to normalize their distribution, then compared via one-way ANOVA with Dunnett’s multiple comparison test.

### Structural similarity search

The highest confidence AlphaFold3 derived models for PilX1 and PilX2 were modified in PyMol 3.0.0^121^ to remove the N-terminal hydrophobic alpha helix and globular pilin domain, leaving just their unique C-terminal domains. The resulting structures were exported as PDBs and submitted to the DALI webserver^72^ for an Exhaustive PDB25 search using default parameters^73^. Outputs were screened for functional categories and those with roles in carbohydrate/glycan hydrolysis or binding were highlighted in green (Supplemental table 3). The first carbohydrate binding protein of bacterial origin from each DALI output (5KLE and 6GXS respectively) was then aligned against the original query structure using the RCSB protein Data Bank alignment tool^122^ and the jFATCAT (rigid) method^123^.

### *S. odontolytica* genetics

*S. odontolytica* XH001 was made electrocompetent and transformed as previously described^124–126^. Briefly, to generate constructs 1,000 bp homology arms flanking *rmlC/D* (APY09_RS00100), *rgpB* (APY09_RS05560), *rgpF* (APY09_RS05510), *tagO* (APY09_RS06605), a putative galactosyltransferase (APY09_RS05455), a putative N-acetyl glucosyltransferase/*gacI* (APY09_RS00090), and a convergently encoded neutral site termed NSXH (between APY09_RS05760 and APY09_RS05765) were amplified and assembled into linear allelic exchange constructs using either the Kanamycin resistance cassette from pJRD215^127^, the chloramphenicol resistance cassette from pJRD215-catP and pCWU6^128^, or the Erythromycin resistance cassette from pVA2198^129^. To generate competent cells, cultures of *S. odontolytica* were grown to midlog phase, treated with 3.75% glycine (w/v) to permeabilize cells, rinsed in water once, and 10% glycerol twice. 1000-2000 ng of linear template was added to competent cells before electroporation at 25 mF, 2.5 kV, and 400 ohms. Selection utilized 400 μg/mL kanamycin, 10 μg/mL thiamphenicol, or 300 μg/mL erythromycin as appropriate.

### Cell wall isolation and modified anthrone assay

Bacterial cell wall purification protocol was based on a peptidoglycan isolation protocol^130^, and modified to exclude glycan removing steps. Briefly, 100-500 mL cultures of each strain were grown anaerobically overnight. Ampicillin (150 μg/mL) was added for 30 minutes. Cells were rinsed, frozen, and then added dropwise into boiling PBS with 8% SDS (w/v) for 30 minutes. Liberated sacculi were rinsed thrice, and then treated with DNase (10 μg/mL), RNase (50 μg/mL), and Pronase (200 μg/mL). All enzymatic digestions performed at 37°C for 90 minutes. Finally, sacculi were frozen, added to 1% PBS and boiled for 30 minutes, rinsed twice in water, rinsed once in 100% acetone, dehydrated, and weighed.

Our modified anthrone assay was developed based on previous analyses of Rha-CWPS in *Streptococcus mutans*^85^. Briefly, a 0.2% (w/v) solution of anthrone in concentrated sulfuric acid was prepared. For each sample, 1 mL of anthrone solution was added to 200 μL of monosaccharides in water (0.1% and 0.01% w/v) or 200 μL of purified sacculi in water (2 mg and 0.2 mg mass). Samples were heated to 98°C for 10 minutes then cooled prior to spectroscopy. Based on the absorption spectra of fucose, rhamnose, and glucose, OD was measured at 580 (representing all carbohydrates present) and 690 (representing the contribution of glucose specifically). Rhamnose/fucose to glucose ratio was calculated as (OD_580_-OD_690_)/OD_690_.

### Saccharibacteria competition

Prior to re-association experiments, all Saccharibacteria strains had to be isolated as previously described and quantified via qPCR with strain-specific primers^50^. All competitions were performed in biological triplicate. For tripartite competition, TM7x, BB004, and TM7-008 were each introduced into overnight *S. odontolytica* cultures with individual MOIs of 0.1, followed by outgrowth and passaging into fresh media (with or without 50 mM L-rhamnose) every 24 hours for 12 days. Pairwise competitions were performed by using established co-cultures of *S. odontolytica* infected with each of the experimental Saccharibacteria, separately infecting each co-culture with the remaining two species at an MOI of 0.1 and passaging every 24 hours for 8-12 days. Mutant TM7x competition was performed using an established co-culture containing BB004 with each TM7x genotype at an MOI of 0.1 and then passaging for 10 days.

### Fluorescence *In Situ* Hybridization and Wheat Germ Agglutinin staining

All oligonucleotide probes are described in supplemental table 5. Whole-mount FISH was performed as previously described^131,132^ with adjustments. Briefly, bacterial competition experiments were set up as described above and at indicated passages, 1 mL of co-cultures were fixed in 1 mL fixation solution (4% formaldehyde in 1 x PBS) at 4°C for 1 hour. Fixed cells were digested with 2mg/mL lysozyme at 37C for 9 minutes. Cells were incubated in 50% ethanol for 5 minutes, then 80% ethanol, then 90% ethanol. Cells were washed and re-suspended in 200 μL hybridization solution (900mM NaCl, 20mM Tris, pH 7.5, 0.01% SDS, 20% formamide) then incubated for 2 hours in hybridization solution supplemented with appropriate probes at a final concentration of 5 ng/μL (0.78 μM) at the indicated temperatures (Table S6).

Keep all staining in the dark after this step. Cell were then washed in wash buffer (215 mM NaCl, 20 mM Tris, pH 7.5, 5 mM EDTA) at 48°C for 15 min, then further washed in two changes of 0.1x SSC (15 mM NaCl, 150 μM sodium citrate) at 37 °C for 10 minutes each. If using WGA staining, during the last wash, add WGA conjugated to Alexa Flour 488 (ex:490/em:520) to wash buffer at a concentration of 50 μg/mL and incubate at room temperature for 30 minutes. Spread bacteria on PolyLys coated cover slips or adherent glass slide (Ultra frost with ultra stick Adhesin, Cat No 3039) and incubate for 10 minutes at room temperature. Wash cells with water, dry, and apply cover slip with 2.5 μL of SlowFade (antifade reagent). Imaging was performed using a Zeiss LSM 780 Confocal Microscope with a 40x 1.4 N.A. Plan-Apochromat objective.

## Authors’ contributions

Conceptualization by ASG, XH, and BB. Methodology by ASG, JMB, Jett Liu, JBSS, RJL, JSM, and BB. Software by JMB, and JSM. Formal analysis by ASG. Investigation by ASG, LL, JMB, NN, Jett Liu, JDSS, and BB. Resources provided by Jun Liu, RJL, JSM, XH, and BB. Data Curation by JSM. Original Draft written by ASG. Review and Editing by ASG, JSM, XH, and BB. Visualization by ASG, JMB, and BB. Supervision by XH and BB. Funding acquisition by JSM, XH, and BB.

## Acknowledgements

Some of the microscopy images were acquired at the ADA Forsyth Institute Advanced Microscopy Core Facility (RRID:SCR_021121). Cryo-ET data were collected at Yale CryoEM Resource, which was funded in part by the NIH grant 1S10OD023603-01A1. Hygromycin B resistance cassette generously provided by Dr. Larry A. Gallagher and Dr. Joseph D. Mougous. We thank Dr. Joseph K. Bedree for helpful discussions on *S. odontolytica* genetics and competency. Thiamphenicol/Chloramphenicol resistance cassette generously provided by Dr. Chenggang Wu. We thank Otari Chipashvili for method development for Saccharibacteria competitions.

---

**Supplemental figure 1.**
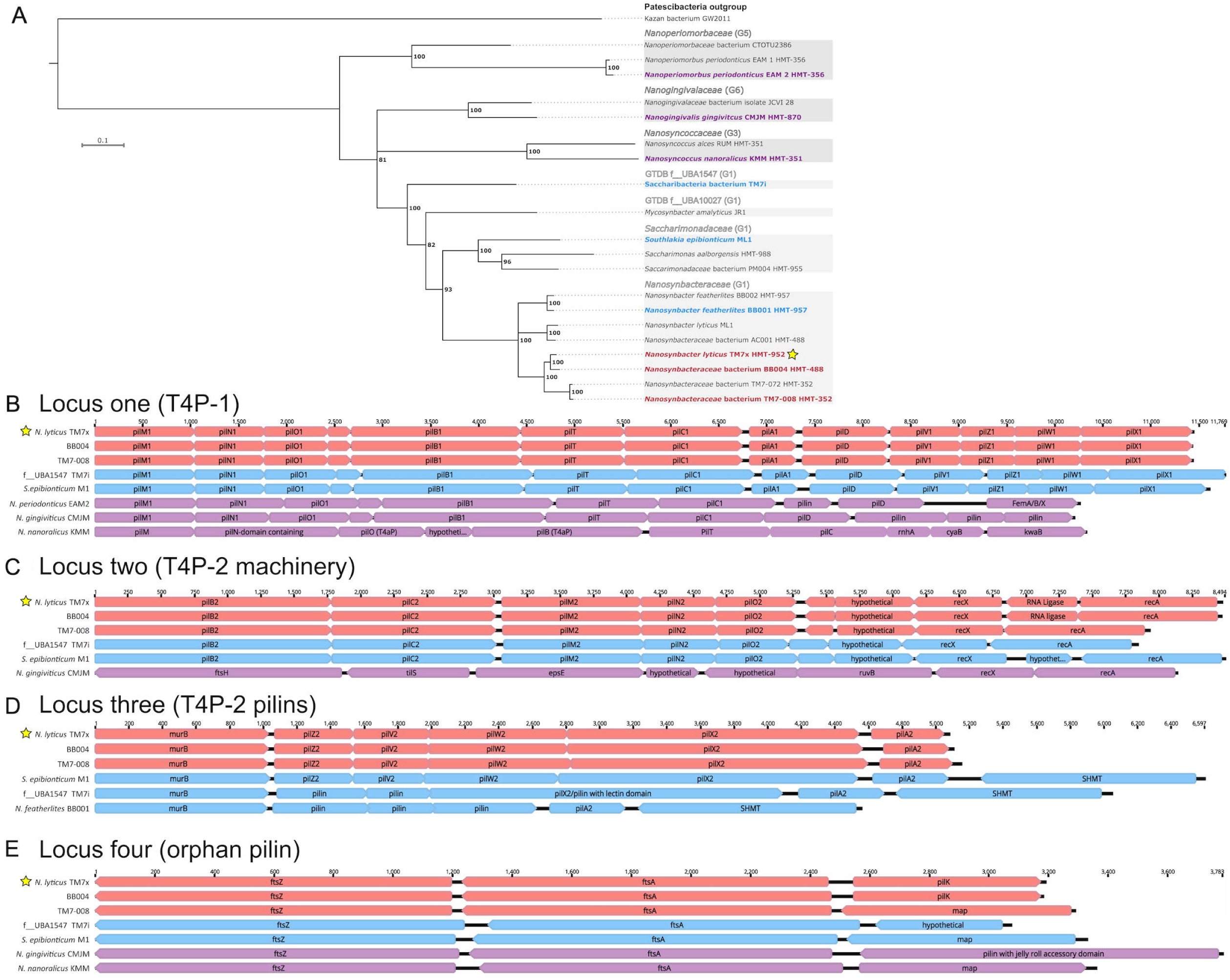
Conservation of T4P from TM7x across the phylum Saccharibacteria. **(A)** a maximum likelihood phylogeny depicting representative Saccharibacteria strains with complete genomes reveals that *N. lyticus* falls within the G1 clade, while more distant relatives cluster into G3, G6, and G5. <u>H</u>uman <u>m</u>icrobial taxon (HMT) IDs provided for strain within eHOMD. For analysis of synteny amongst T4P loci, a subset of genomes present on the phylogeny were compared. Close relatives of TM7x are indicated in red, intermediate relatives are indicated in blue, and distant relatives are indicated in purple. **(B)** examination of locus one indicates that T4P-1 is present in all examined sequences, however it has become highly divergent in TM7-KMM. Examination of locus two **(C)** and three **(D)** indicate strong conservation of T4P-2 amongst all G1 Saccharibacteria, and complete absence outside of G1. **(E)** The orphan pilin at locus 4 is seemingly specific to TM7x and BB004, however of note, the distantly related TM7-CMJM had a completely different putative minor pilin present at this locus.

**Supplemental figure 2.**
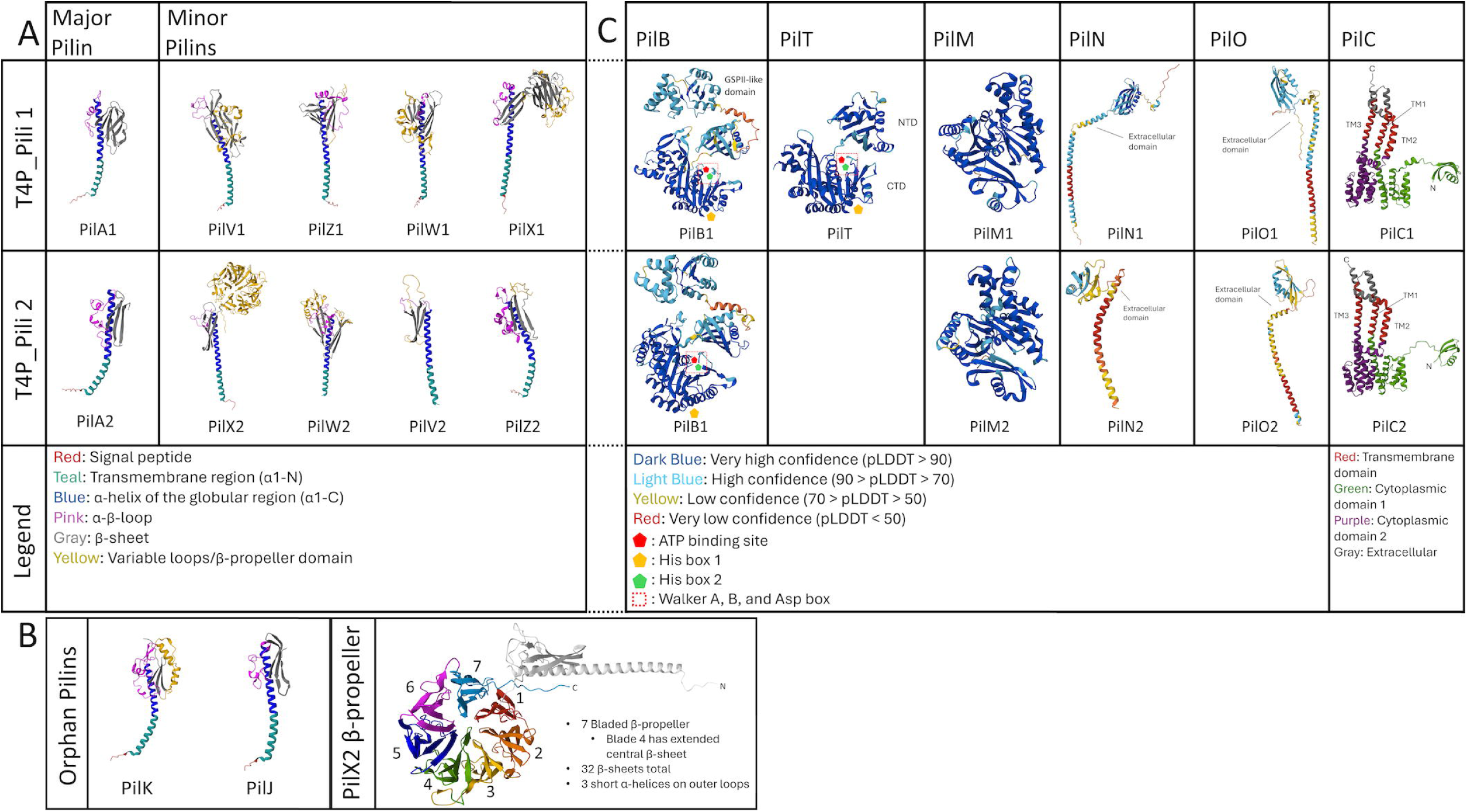
AlphaFold2 structural models of T4P proteins from TM7x. Structural models of T4P pilins **(A-B)** and assembly machinery **(C)** encoded in all four identified loci, comparing the essential T4P-1 system (row one) and the non-essential T4P-2 (row two), notably T4P-2 has no associated retraction ATPase/PilT. The legend (row three) indicates either the model confidence level reported by AlphaFold2 (PilBTMNO) or indicates specific functional structures (PilC and the pilin proteins). Special attention is given to the orphan pilins which cannot be confidently assigned to either T4P system as well as the relatively unique PilX2 structure **(B)**.

**Supplemental figure 3.**
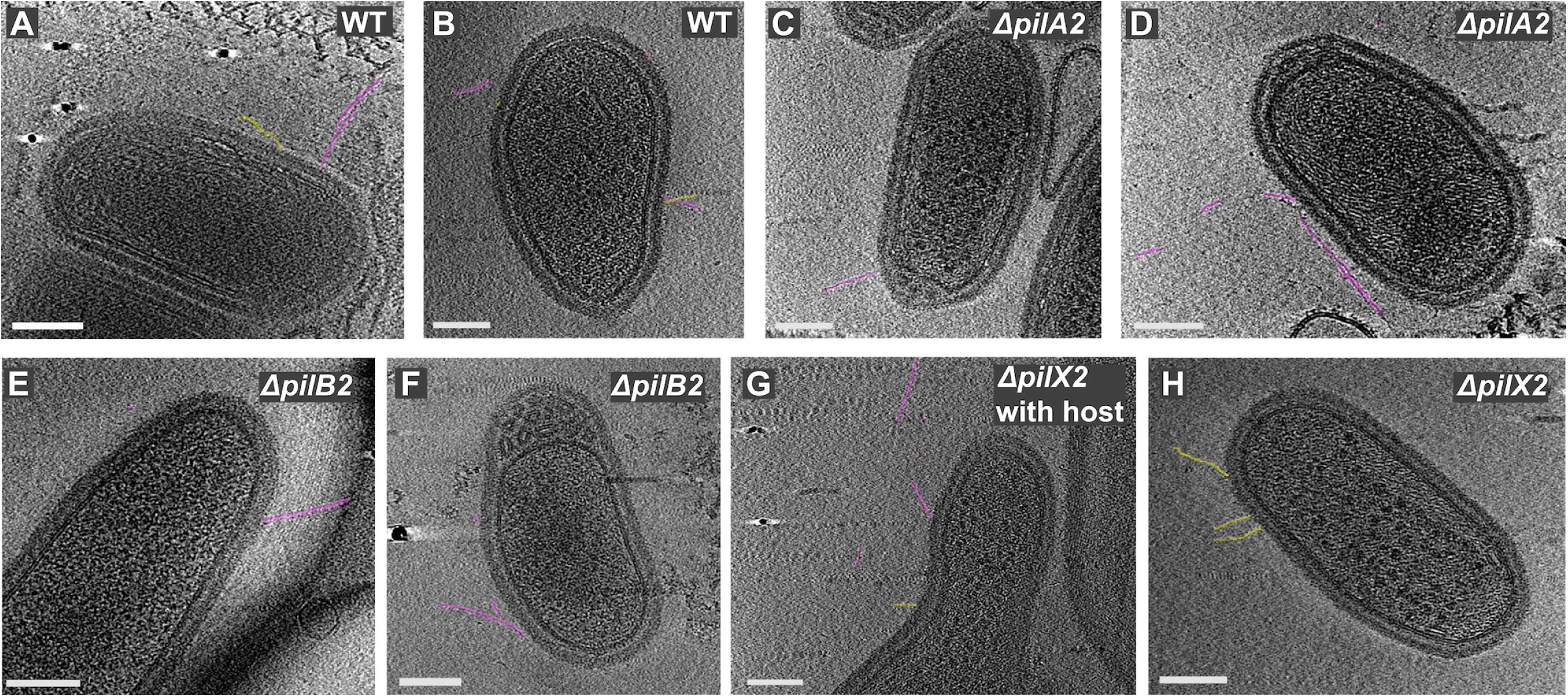
Expanded Cryo-ET images. Additional Electron tomography images of *N. lyticus* TM7x wildtype **(A-B),** TM7xΔ*pilA2* **(C-D)**, TM7xΔ*pilB2* **(E-F)**, and TM7xΔ*pilX2* **(G-H)**. All scale bars are 100 nm. Select pili filaments are highlighted to indicate either thin pili (yellow; diameter ≈1.8 nm) or thick pili (purple; diameter ≈3.2 nm). Thin filaments are only detectable in wildtype cells and TM7xΔ*pilB2*, thick filaments are seen in all treatments. All images show planktonic cells except panel G which shows a cell near, but not in close contact with, a host bacterium (host on right edge).

**Supplemental figure 4.**
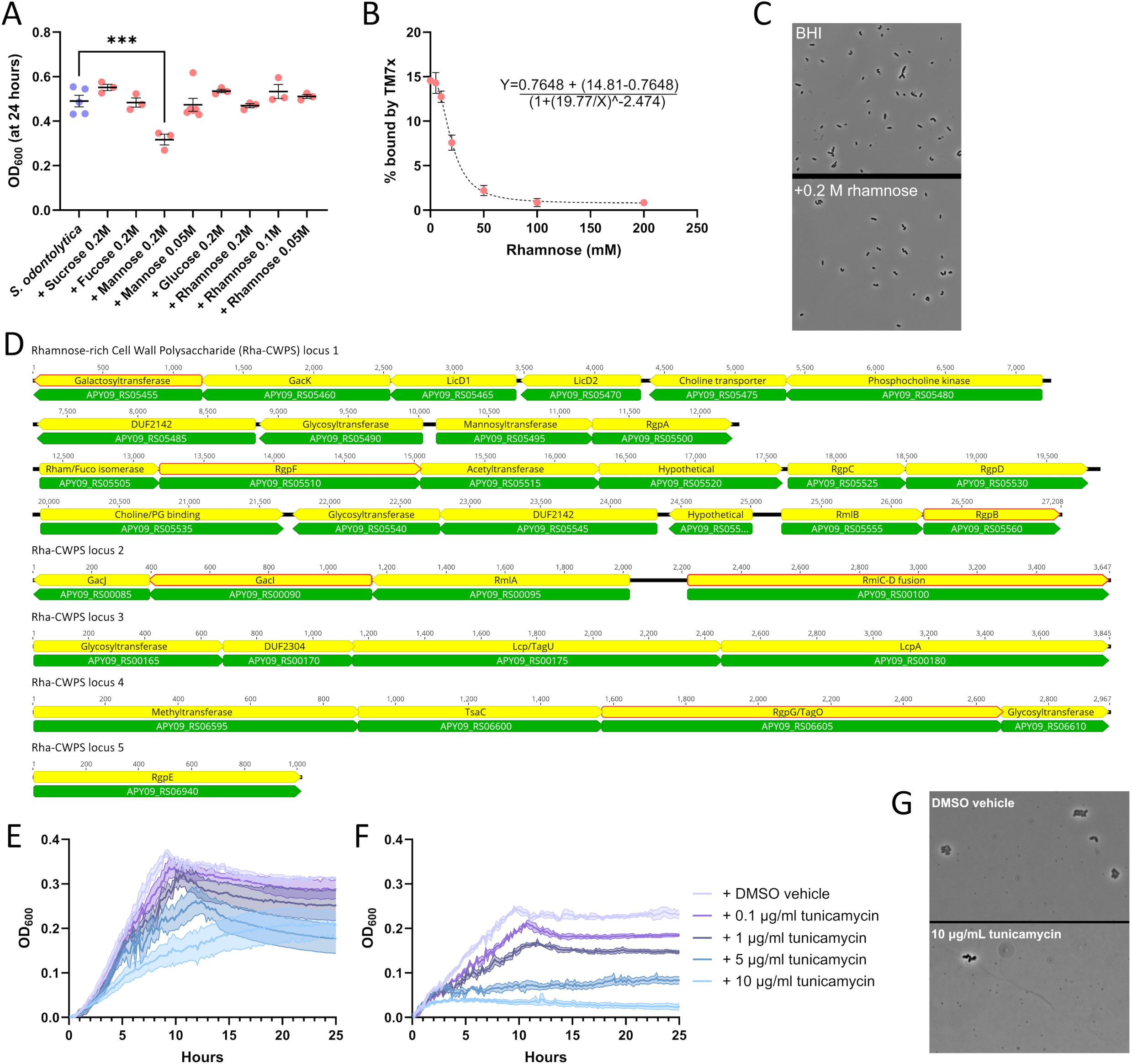
Alignment to carbohydrate binding proteins, Rha-CWPS biosynthesis loci, and tunicamycin treatment. *S. odontolytica* was grown in the presence of diverse monosaccaride sugars before testing TM7x binding **(A)**, and only high concentrations of mannose negatively impacted growth. The monosaccharide rhamnose caused a dose dependent inhibition of TM7x binding **(B)**, here modeled with a 4-parameter non-linear fit (R^2^ = 0.975). **(C)** Representative images show the inhibition of TM7x binding. S*. odontolytica* encodes homologs of rhamnose-rich cell wall polysaccharide (Rha-CWPS) synthesis genes **(D)**. These included genes for UDP-Rhamnose synthesis, rhamnose extension, cell wall attachment, phosphocholine decoration, and glycosyltransferase decoration. Allelic replacement was attempted for 6 genes (indicated in red) however all except for the *gacI* homolog (APY09_RS00090) were essential. Tunicamycin, a chemical inhibitor of TagO (aka RgpG) was applied to *S. odontolytica* **(E)** as well as TM7x containing co-culture **(F)** to optimize inhibition. Tunicamycin appeared to inhibit host growth far more than TM7x growth **(G)**. * = p ≤ 0.05, ** = p ≤ 0.01, *** = p ≤ 0.001, **** = p ≤ 0.0001.

**Supplemental figure 5.**
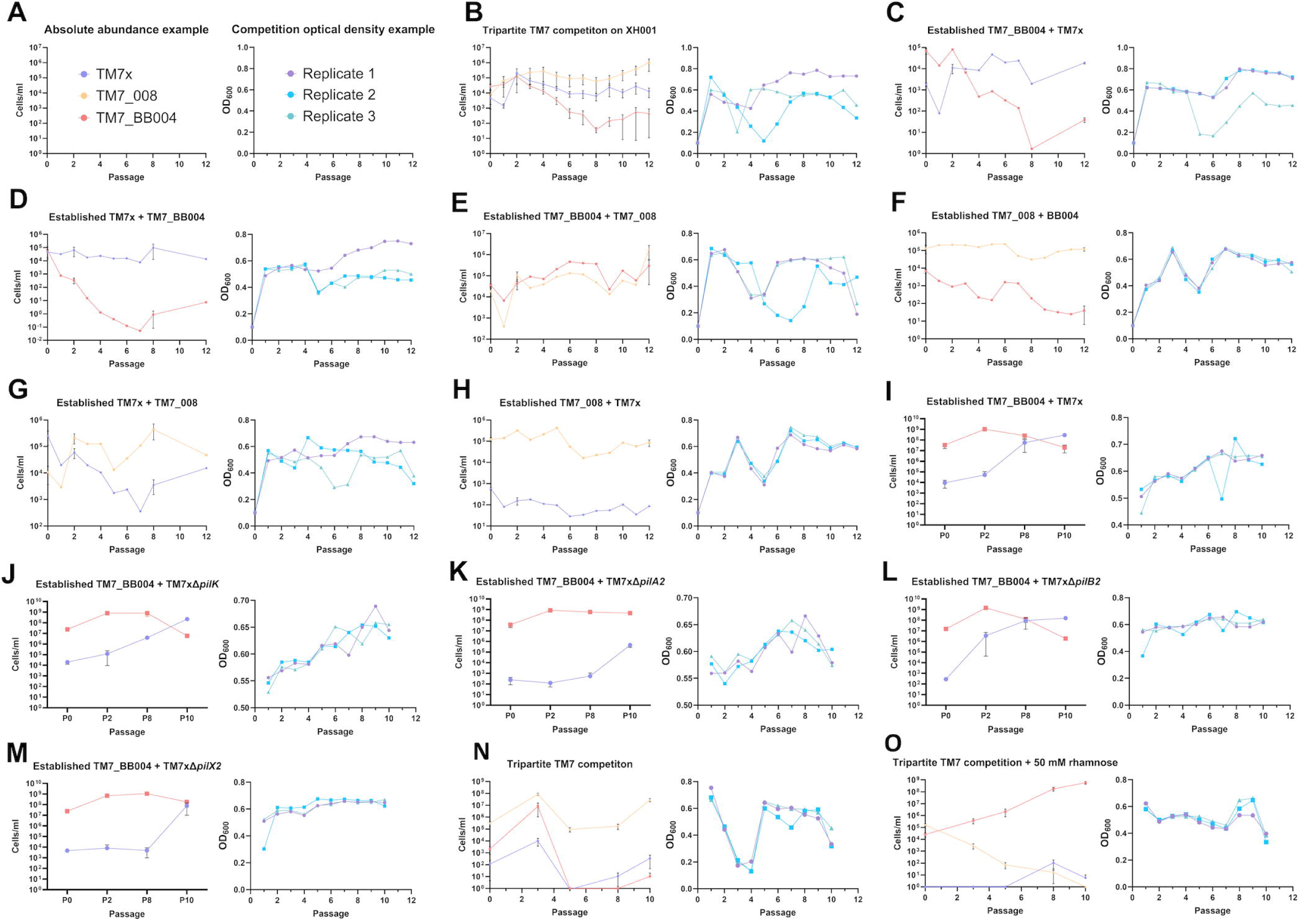
Absolute abundance and OD_600_ data from all competition experiments. Paired qPCR enumeration of TM7x, TM7-008, and BB004 and co-culture optical density values for all competition experiments performed, with universal key in panel **A**. Tripartite competition **(B)** indicates a clear and repeatable outcome where BB004 declines, TM7x persists, and TM7-008 grows to dominance. This is borne out by pairwise combinations of TM7x with BB004 **(C-D)** which indicate that TM7x has higher competitive fitness on this host. Pairwise combinations of BB004 with TM7-008 **(E-F)** and TM7x with TM7-008 **(G-H)** similarly support the ranked competitive fitness observed in the tripartite competition. To examine potential priority effects, each competition was performed twice, each time starting with a different competitor in established co-culture with the host. Competition of BB004 with wildtype TM7x **(I)**, TM7xΔ*pilK* **(J)**, TM7xΔ*pilA2* **(K)**, TM7xΔ*pilB2* **(L)**, and TM7xΔ*pilX2* **(M)** shows that some pilin mutants have decreased competitive fitness.

**Supplemental figure 6.**
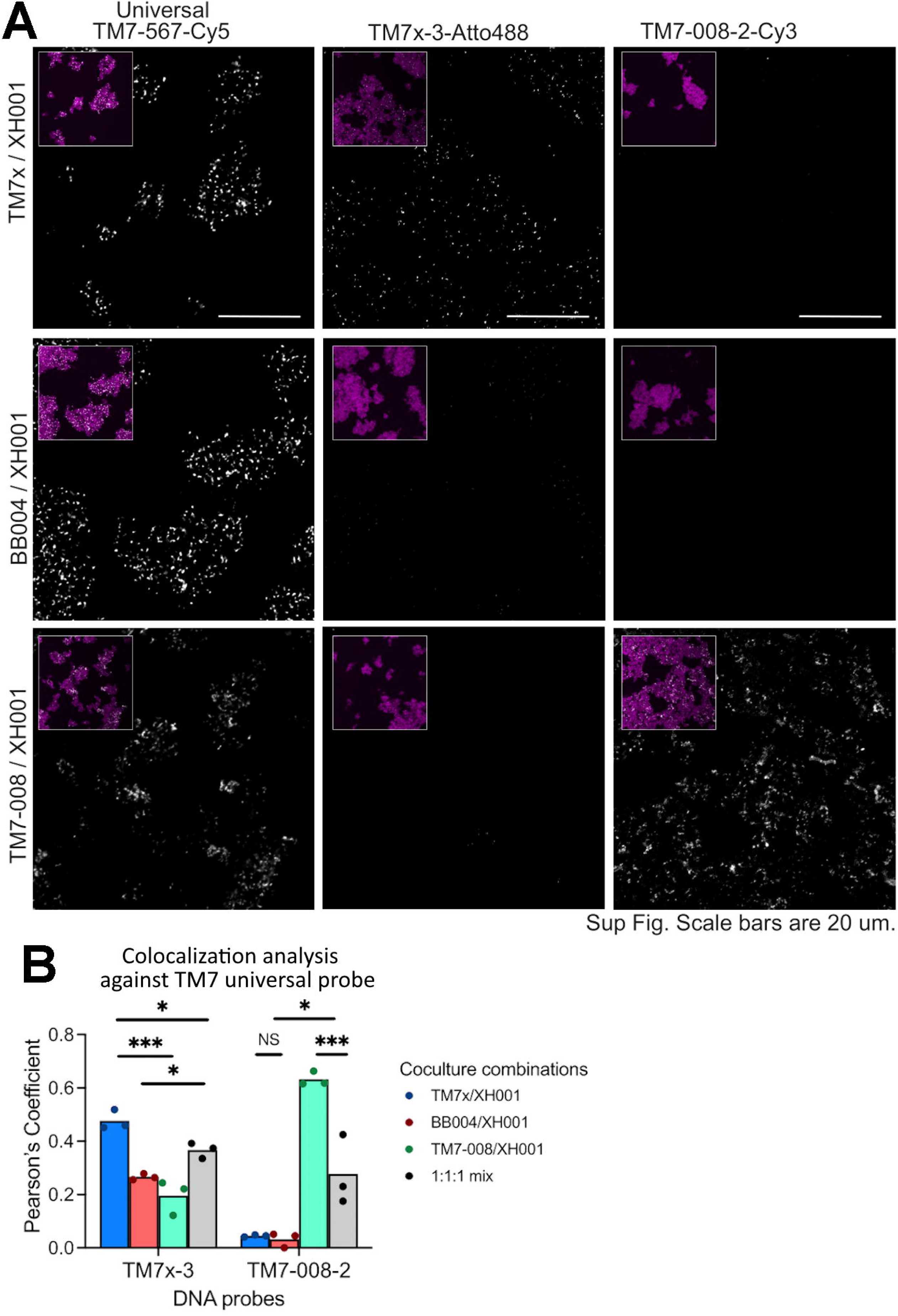
Optimization of strain-specific Saccharibacteria FISH probes. To ascertain the specificity of 16s oligonucleotide FISH probes, each probe was conjugated to a distinct fluorophore and used to stain fixed monoxenic co-cultures of *S. odontolytica* containing TM7x, BB004, and TM7-008 **(A)**. To quantify these specificities, cultures were stained with both a strain-specific probe and a universal Saccharibacteria probe (TM7-567), then colocalization of both probes, as approximated via the Pearsons coefficient, was used to quantify the binding potential of each probe for each strain **(B)**. Due to the similarity of 16s sequences, no tested probes were highly specific for BB004, however TM7x-3 displayed some specificity for TM7x, and TM7-008-2 displayed a high level of specificity for TM7-008. * = p ≤ 0.05, ** = p ≤ 0.01, *** = p ≤ 0.001, **** = p ≤ 0.0001.

**Supplemental figure 7.**
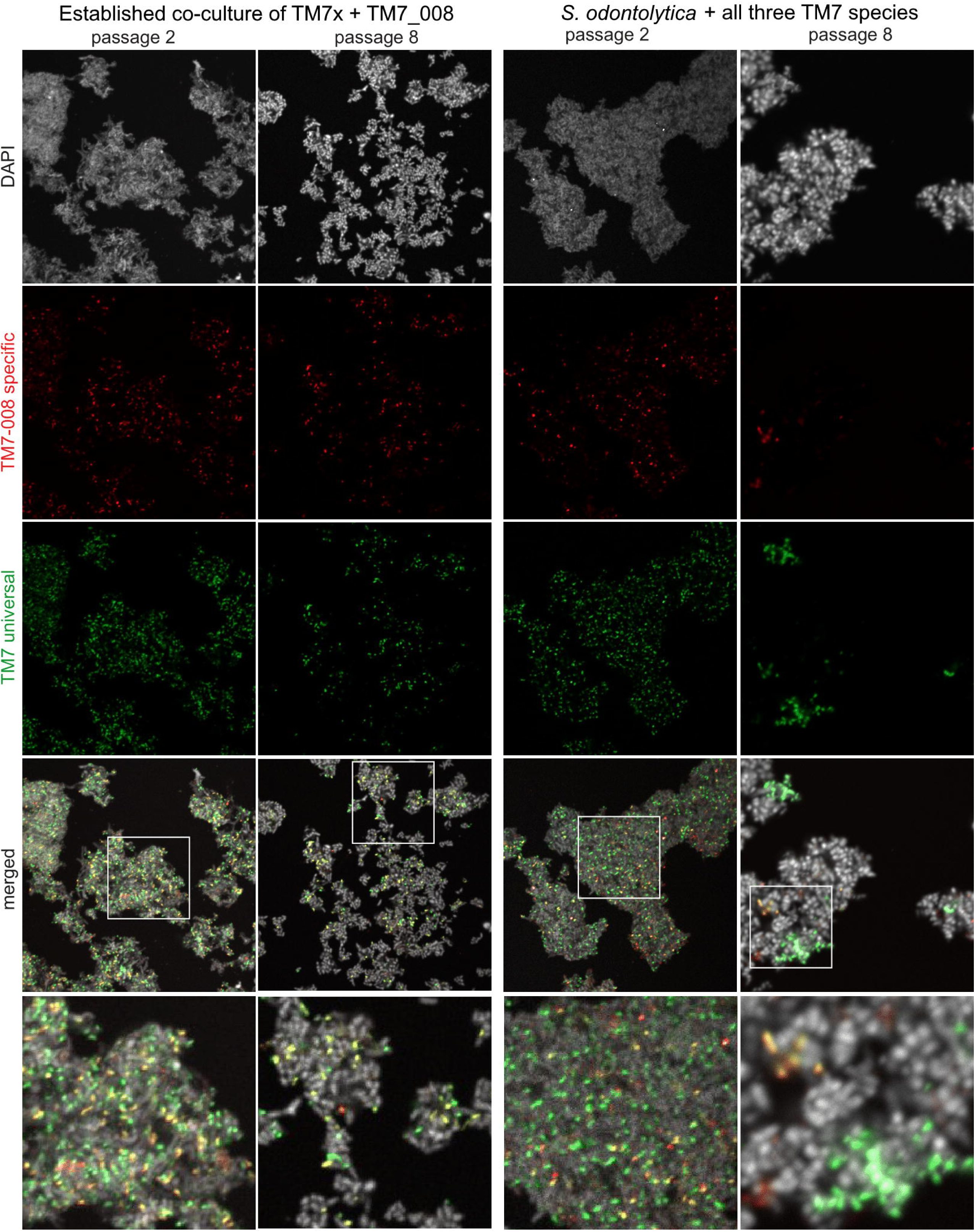
Spatial visualization of Saccharibacteria competition cultures. To visualize the fine scale arrangement of competition between Saccharibacteria strains for a single bacterial host, fixed co-cultures were stained with DAPI (host cells shown in white), TM7-008-2 probes (TM7-008 specific shown in red), and TM7-567 probes (universal TM7 shown in green). Two distinct competition conditions, established TM7x infected with TM7-008 and tripartite infection, were visualized at two different times, passage two and passage eight post infection. At passage two, competitors were evenly distributed throughout the clumped biomass of actinobacteria with TM7-008 appearing as red or yellow puncti while all other Saccharibacteria appear green. Passage eight cultures show more organized arrangement of episymbionts, particularly in the tripartite co-cultures, where distinct strain level clusters are visible amongst the mass of bacterial host cells.

## Supplemental materials

Supplemental table 1: Similarity based *pil* gene annotation

Supplemental table 2: Patescibacteria database metadata

Supplemental table 3: DALI structural similarity search outputs

Supplemental table 4: Compiled list of all primers used in this study

Supplemental table 5: Compiled list of all fluorescent dyes/FISH probes used in this manuscript and their information (Company, Ex/Em, etc.)

Supplemental table 6: Compiled list of all strains and their growth conditions

